# Cardiac sympathovagal activity initiates a functional brain-body response to emotional processing

**DOI:** 10.1101/2021.06.05.447188

**Authors:** Diego Candia-Rivera, Vincenzo Catrambone, Julian F. Thayer, Claudio Gentili, Gaetano Valenza

## Abstract

A century-long debate on bodily states and emotions persists. While the involvement of bodily activity in emotion physiology is widely recognized, the specificity and causal role of such activity related to brain dynamics has not yet been demonstrated. We hypothesize that the peripheral neural monitoring and control of cardiovascular activity prompts and sustains brain dynamics during an emotional experience, so these afferent inputs are processed by the brain by triggering a concurrent efferent information transfer to the body. To this end, we investigated the functional brain-heart interplay under emotion elicitation in publicly available data from 62 healthy participants using a computational model based on synthetic data generation of EEG and ECG signals. Our findings show that sympathovagal activity plays a leading and causal role in initiating the emotional response, in which ascending modulations from vagal activity precede neural dynamics and correlate to the reported level of arousal. The subsequent dynamic interplay observed between the central and autonomic nervous systems sustains emotional processing. These findings should be particularly revealing for the psychophysiology and neuroscience of emotions.

**Significance:** We investigate the temporal dynamics of brain and cardiac activities in healthy subjects who underwent an emotional elicitation through videos. We demonstrate that, within the first few seconds, emotional stimuli modulate the heart activity, which in turn stimulate an emotion-specific cortical response in the brain. Then, the conscious emotional experience is sustained by a bidirectional brain-heart interplay and information exchange. Moreover, the perceived intensity of an emotional stimulus is predicted by the intensity of neural control regulating the heart activity. These findings may constitute the fundamental knowledge linking neurophysiology and psychiatric disorders, including the link between depressive symptoms and cardiovascular disorders.

## Introduction

The role of bodily activity in emotions is often questioned. Despite the vast literature showing bodily correlates with emotions, a long-lasting debate about the relationship between bodily states and emotions persists.^1^The central nervous system (CNS) communicates with the autonomic nervous system (ANS) through interoceptive neural circuits that contribute to physiological functions beyond homeostatic control, including emotional experience.^2^The debate about the role of the ANS in emotions can be condensed into two views: specificity or causation.^1^The specificity view is related to the James–Lange theory, which states that bodily responses precede emotions’ central processing, meaning that bodily states would be a response to the environment, followed by an interpretation carried by the CNS that would result in the emotion felt. However, causation theories represent an updated view of the James–Lange theory, suggesting that afferent autonomic signals from the body influence perceptual activity in the brain.^3,4^In this direction, subjective experiences may be influenced or shaped by ascending communication from visceral inputs.^5–7^

Functional models of CNS and ANS interplay have described bidirectional dynamics in emotions.^8–10^In particular, the functional brain-heart interplay (BHI) involves brain structures that comprise the central autonomic network (CAN), which has been described as being in charge of autonomic control,^11,12^indicating that the CAN primarily performs descending modulations to bodily activity. In addition, the default mode network (DMN) is usually associated with passive states.^13^ Nevertheless, the DMN is also involved in autonomic control^14^ and tasks of self-related cognition and interoception,^15,16^ suggesting that the DMN participates in both ascending and descending communications with the heart. The evidence on the role of the heart in subjective experiences spans from perception, self-related cognition, to consciousness.^7^

Functional neuroimaging studies have uncovered several correlates of different autonomic signals in the brain during emotional experiences.^17–19^To understand these correlations and the functional interactions between the heart and brain, various methods have been proposed to investigate functional BHI through non-invasive recordings.^20^The study of emotions using these methods comprises the analysis of heartbeat-evoked potentials,^21^non-linear couplings,^22^and information transfer modeling.^23^However, the role of bodily input remains unknown.^1^

Advances in physiological modeling have contributed considerably to the understanding of brain-body interactions and emotions.^5,8,10,24^In this study, we take a step forward and investigate through a mathematical model-based approach whether such peripheral neural dynamics play a causal role in the genesis of emotions. We applied a model of BHI based on synthetic data generation (SDG),^25^estimating the directionality of the functional interplay using simultaneous electroencephalography (EEG) and electrocardiography (ECG) recordings gathered from healthy subjects undergoing emotion elicitations with video clips, the publicly-available *DEAP* and *MAHNOB* datasets.^26,27^We hypothesize that emotion elicitation follows causal interactions with the ANS, in which the CNS integrates the afferent information flow, namely from-heart-to-brain interplay, which generates a cascade of neural activations that, in turn, modulate directed neural control onto the heart, namely from-brain-to-heart interplay.

## Materials and methods

### 1. Datasets description

This study comprised the analysis of male and female healthy human volunteers from two publicly available datasets undergoing video stimulations with affective content and physiological signals acquisition.

#### Dataset I

The *DEAP dataset for emotion analysis*^26^ is available at http://www.eecs.qmul.ac.uk/mmv/datasets/deap/. The dataset consisted of 32 subjects who underwent video visualizations. Data were collected at 512 Hz using 32-channel EEG and 3-lead ECG. Additional physiological signals were not considered in this study. Data from all participants were used (age range, 19–27 years; median, 27 years; 16 females).

The dataset consisted of 40 video trials from music videos (see the Appendix for details). Videos had a duration of 60 s and were presented after an initial resting period of 120 s. Trials have a pad of 5 s at the beginning and 3 s at the end. In this study, the trials were compared to the average initial 120 s rest period.

#### Dataset II

The *MAHNOB–HCI dataset of emotion elicitation*^27^ is available at https://mahnob-db.eu/hci-tagging/. The dataset consisted of 30 subjects who underwent video visualizations. Data were collected at 256 Hz using 32-channel EEG and 3-lead ECG. Additional physiological signals were not considered in this study. A total of 27 subjects participated in the study (age range, 19–40 years; median, 26 years; 15 females). Data from individual trials involved a different number of participants, which ranged between and 25–27, either because physiological data were not available at the moment (accessed May 7^th^, 2020), or the quality of their ECG was not sufficient to properly identify R-peaks. The dataset consists of 20 video trials from movies (see the Appendix for details). Videos had a duration of 35–117 s and were presented between neutral videos of approximately 20 s duration. Trials had a pad of 30 s at the beginning and end. In this study, the trials were compared to the average 30 s rest period before each trial.

Subjective ratings of the emotional experience in the two datasets rely on the circumplex model of affect, which considers a two-dimensional approach to classify emotions: valence related to pleasantness and arousal related to intensity. In this view, emotions can be determined by a linear combination of these two dimensions. We subdivided the trials of the datasets into three groups of emotions, based on group median valence and arousal from the self-assessment values: pleasant (high valence and high arousal), unpleasant (low valence and high arousal), and neutral (unrelated to valence and low arousal). The thresholds of valence and arousal were selected independently for both datasets to obtain an equal as possible number of trials in each of the three subgroups (see Appendix for details).

### 1. EEG processing

The aim of the EEG pre-processing was to obtain a clean and artifact-free EEG to consecutively compute the EEG spectrogram. The entire process involves frequency filtering, large artifact removal, eye movements, cardiac-field artifact removal, and interpolation of contaminated channels. The process was performed using MATLAB R2018b (MathWorks) and Fieldtrip Toolbox.^28^EEG data were bandpass filtered with a Butterworth filter of order 4, between 0.5 and 45 Hz. Large artifacts were removed using wavelet-enhanced independent component analysis (Wavelet-ICA),^29^which were identified using automated thresholding over the independent component and multiplied by ×50 to remove only very large artifacts. EEG data were reconstructed, and ICA was re-run to identify eye movements and cardiac-field artifacts from the EEG data. The process involved one lead from the ECG as an additional input in the ICA to ease the process of finding cardiac artifacts. Once the ICA components with eye movements and cardiac artifacts were visually identified, they were set to zero to reconstruct the EEG series. Individual channels were examined under two criteria in the case of noise remnants in the EEG data. Channels were marked as contaminated if their area under the curve exceeded three standard deviations of the mean of all channels. The remaining channels were compared with their weighted-by-distance-correlation neighbors using the standard Fieldtrip neighbor’s definition for 32 channels Biosemi system. If a channel resulted in a weighted-by-distance correlation coefficient of less than 0.5, it was considered contaminated. A maximum of three channels were discarded per recording by the first criterion, and a maximum of six altogether using the two criteria. After analyzing the EEG channels under the second criterion, more than six channels were marked as contaminated; only the channels with lower correlation coefficients with the neighbors were discarded until only six channels were discarded. The contaminated channels were replaced by the neighbor’s interpolation, as implemented in Fieldtrip. Channels were re-referenced offline using a common average.^20^

The EEG spectrogram was computed using a short-time Fourier transform with a Hanning taper. The calculations were performed with a sliding time window of 2 s with a 50% overlap, resulting in a spectrogram resolution of 1 s and 0.5 Hz. Successively, time series were integrated within five frequency bands: delta (δ; 0–4 Hz), theta (θ; 4–8 Hz), alpha (α; 8–12 Hz), beta (β; 12–*f*30 Hz) and gamma (γ; 30–45 Hz).

### 2. ECG processing

The goal of the ECG pre-processing was to obtain R-peak timing occurrences and consecutively compute low-and high-frequency heart rate variability components. The whole process involves frequency filtering, R-peak detection, correction of misdetections, and determination of whether the recording is of optimal quality to be considered in the study. ECG data were bandpass filtered using a Butterworth filter of order 4, between 0.5 and 45 Hz. Heartbeats from QRS waves were identified in an automated process based on a template-based method for detecting R-peaks.^20^Misdetections were corrected first by visual inspection of detected peaks and the respective inter-beat interval histogram, and then automatically using a point-process algorithm.^30^Recordings presenting segments with unintelligible R-peaks were disqualified from the analysis.

The heart-rate variability (HRV) series were studied in the low-frequency (LF: 0.04–0.15 Hz) and high-frequency (0.15–0.4 Hz) ranges to quantify the sympathovagal and parasympathetic activity from the ANS, respectively. Once the heartbeats were detected from the ECG, the HRV series were constructed as the inter-beat interval duration time course. Consecutively, the HRV series were evenly resampled at 4 Hz using spline interpolation. The HRV power was computed using a smoothed pseudo-Wigner–Ville distribution.^31^The pseudo-Wigner–Ville algorithm consists of a two-dimensional Fourier transform with an ambiguity function kernel to perform two-dimensional filtering. The ambiguity function comprises ellipses whose eccentricities depend on the parameters *ν0* and *τ0*, setting the filtering degrees of the time and frequency domains, respectively, and an additional parameter *λ* was set to control the frequency filter roll-off and kernel tail size.^31^The following parameters were considered in this study: *v0* = 0.03, *τ0* = 0.06, and *λ* = 0.3. ^25,31^

### 3. Computation of directional, functional brain–heart interplay

Functional brain-heart interplay was assessed using the synthetic data generation (SDG) model,^25^whose software implementation is publicly available at https://github.com/CatramboneVincenzo/Brain-Heart-Interaction-Indexes. The model provides a time course of the estimated coupling coefficients between the different heart and brain components studied separately for both possible modulation directions.

#### Functional Interplay from the brain to the heart

This was quantified through a model able to generate synthetic heartbeat intervals based on an integral pulse frequency modulation model, which is parameterized with Poincaré plot features.^32^The synthetic heartbeats were modeled as Dirac functions *δ(t)*, generating an impulse at the timings of heartbeat occurrences t*_k_*, as presented in Eq. (1). Heartbeat generation is triggered by the integral of a reference heart rate μ*_HR_* and a modulation function *m*(*t*), as shown in Eq. (2), where a new R-peak is generated when the integral function reaches a threshold value of 1.

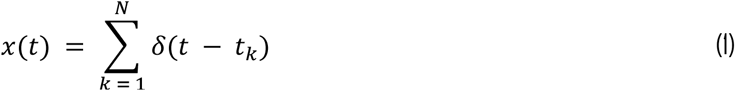

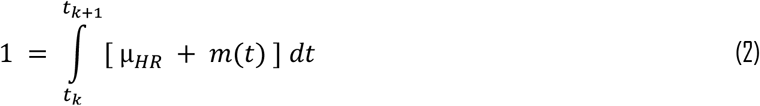

The modulation function *m*(*t*) is represented as a summation of two oscillators on behalf of the sympathetic and parasympathetic autonomic outflows, as presented in Eq. (3). The oscillators are centered at the frequencies ω*_s_* and ω*_p_*, with amplitudes defined by C_S_ and C_P_ indicating time-varying coupling constants, representing the sympathetic and parasympathetic activities, respectively. The coupling constants are defined in Eqs. (4) and (5), where the parameters *L* and *W* are the length and width of the Poincaré Plot,^32^ and γ = *sin*(ω*_p_* / 2μ*_HR_*) − *sin*(ω*_s_* / 2μ*_HR_*).

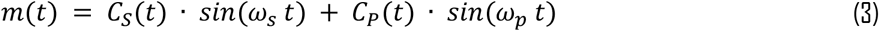

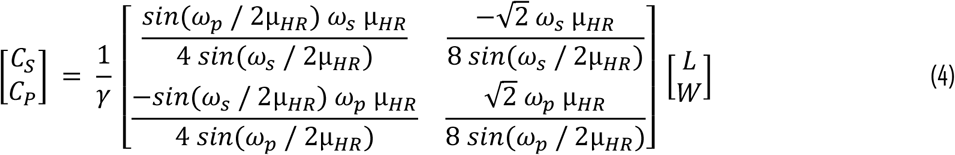

The model considers the interaction from the CNS as the ratio between the coupling constants (i.e., C*_S_* and C*_P_*) and the EEG power in the previous time window Power_f_(*t* − 1) at a defined frequency band *f* (i.e., δ, θ, α, β, and γ). Therefore, the brain-heart interplay coefficients SDG_brain_f →LF_ and SDG_brain_f →-HF_ are defined by Eqs. (5) and (6), respectively.

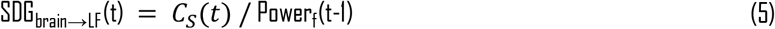

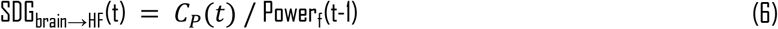

#### Functional Interplay from the heart to the brain

The functional interplay from the heart to the brain is quantified through a model based on the generation of synthetic EEG series using an adaptive Markov process,^33^as shown in Eq. (7). The model estimates the modulations to the brain expressed by the coefficient *Ψ_f_* using least squares in an exogenous autoregressive process, as shown in Eq. (8), where *f* is the main frequency in a defined frequency band, *θ_f_* is the phase, *κ_f_* is a constant, and *ε_f_* is a Gaussian white noise term.

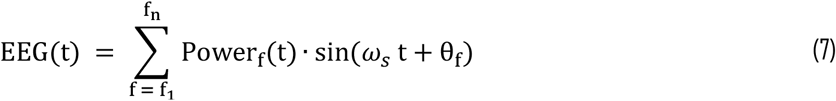

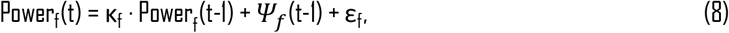

Therefore, the Markovian neural activity generation within a specific EEG channel and frequency band uses its previous neural activity and heartbeat dynamics as inputs for EEG data generation. The coupling coefficients SDG_heart→brain_ can be derived from the contribution of heartbeat dynamics HRV_X_ (with X as LF, HF, or their combination) and the exogenous term of the autoregressive model:

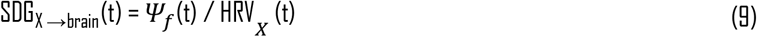

For this study, the model was simulated using a 15 s long time window with a 1 s step to estimate the coefficients. The central frequencies used were ω_s_ = 2Π · 0.1 rad/s and ω*_p_* = 2Π · 0.25 rad/s, in which 0.1 Hz and 0.25 Hz correspond to LF and HF band central frequencies, respectively.

### 4. Statistical analysis

The samples were described group-wise using the median, and the related dispersion measures were expressed as the median absolute deviation. Statistical analyses included the Spearman correlation coefficient, Friedman tests, and cluster-based permutation test, as specified in the main text. Spearman correlation was performed group-wise over individual channels to determine the relationships between the group-median brain-heart coupling coefficients and associated valence or arousal. Correlation coefficient p-values were derived using a Student t-distribution approximation. Friedman tests were performed to assess coupling coefficient changes in their distribution in individual channels, on different time windows, and considering different types of trials (pleasant, unpleasant, neutral). The significance level of the p-values was corrected per the Bonferroni rule, considering the multiple comparisons performed for the 32 EEG channels, with an uncorrected statistical significance set to α = 0.05.

Cluster-based permutation tests^34^were performed over the coupling coefficients between the averaged rest period and the trial of emotion elicitation in their total duration. The comparisons were performed for LF→ brain, HF→ brain, brain →LF, and brain →HF indexes, where the brain part comprises 32 EEG channels, the time course, and the 5 frequency bands studied. The non-parametric cluster-based permutation tests include a preliminary mask definition, identification of candidate clusters, and computation of cluster statistics with Monte Carlo p-value correction. The preliminary mask was computed by performing a paired Wilcoxon test for individual samples defined in space, time, and frequency. If a sample obtained a p-value lower than α = 0.01, the sample was considered part of the preliminary mask. Candidate clusters were formed first on individual timestamps and separately for each frequency band. The identification of neighbor channels was based on the default channels’ neighborhood definition for a 32 channels Biosemi system in the Fieldtrip Toolbox. A minimum cluster size of the three channels was imposed. Adjacent candidate clusters on time were wrapped if they had at least one common channel. Adjacent candidate clusters on frequency were wrapped if they had at least one pair (channel, timestamp) in common. The overall minimum duration of the cluster was set at 2 s.

Cluster statistics were computed from 10,000 random partitions.^34^The data points defined by the cluster’s mask in the space, time, and frequency dimensions were averaged. The samples of the two experimental conditions (rest and emotion elicitation) were placed in a single set. Therefore, samples from the two conditions were randomly selected, and a non-parametric Wilcoxon test was performed over the random partition. The proportion of random partitions that resulted in a lower p-value than the observed one was considered as the Monte Carlo p-value. The cluster’s Monte Carlo p-value was considered to be significant at α = 0.01. The cluster statistic considered is the Wilcoxon’s Z-value obtained from the test on the averaged data points defined by the mask, and the resulting p-value is reported as well. If more than one, the cluster with the highest absolute Z-value was considered.

## Results

### 1. Resting state vs. Emotion elicitation

We computed the SDG model coefficients for individual EEG channels for the entire trial duration, in both the ascending and descending pathways. The output of the model was the time course of the coupling for all combinations of brain oscillations (delta, theta, alpha, beta, and gamma) and heart rate variability, low frequency (LF), and high frequency (HF) components. We investigated the changes in heart-to-brain or brain-to-heart interplay of the resting state with a non-parametric cluster-based permutation analysis. Figure 1 shows the polar histograms counting the number of trials among the two datasets in which the SDG coefficients presented a significant change compared to the resting state. In the *DEAP* dataset, we found that 39 out of the 40 trials presented a significant cluster in either the ascending or descending pathways, whereas in the *MAHNOB* dataset, 17 out of 20 trials. We found that HF-to-brain modulations are more sensitive markers of emotion elicitation than LF-to-brain modulations, whereas brain-to-HF coefficients were more sensitive than brain-to-LF ones. It can be observed that ascending sympathovagal activity, related to the heart rate’s LF component, is usually associated with theta, alpha, and beta oscillations. The ascending inputs from the HF relay to all EEG bands, being slightly less coupled to gamma oscillations. Regarding the brain-to-heart interplay, the LF band is considerably less coupled, with only a few trials in which a significant change was observed. However, modulations over the HF band are presented from all EEG frequency bands, indicating that the HF band is actively coupled with brain dynamics for both ascending and descending pathways. Since significant changes in BHI for emotion elicitations are associated mainly with vagal activity, for the sake of conciseness, in what follows, we focus on results related to HF power (results including the LF power are reported in the Supplementary Material).

**Figure 1.**
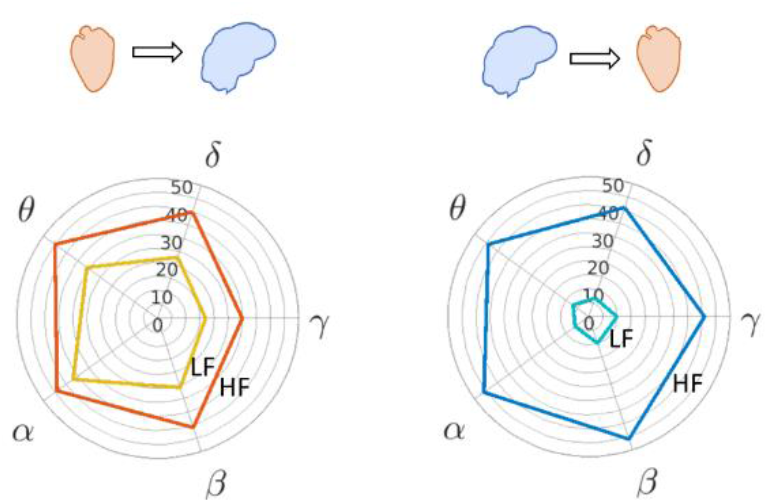
Polar histograms of multiple trials among the two datasets (60 trials) in which the respective heart-to-brain and brain-to-heart components presented a significant change in the cluster-based permutation analysis, with respect to the resting state. The radial-axis indicates the trials’ count, and the angle-axis indicates the EEG frequency band. The left panel presents heart-to-brain interplay. The right panel presents brain-to-heart modulations (See additional material to refer to the polar histograms separately for pleasant, unpleasant, and neutral trials).

After the cluster-based permutation analysis, we combined the segments in which the trials presented a significant change in relation to the resting state (minimum cluster size 3 channels and Monte Carlo p < 0.01) to investigate the scalp regions in which major BHI changes were observed. In Figure 2, we present the SDG coupling coefficient group medians of the HF-to-brain and brain-to-HF values separately for each EEG frequency band. For each subject, SDG coefficients were averaged over time in the significant interval identified by the cluster-based permutation analysis, group medians were computed per trial, and the median among trials was computed afterward. Scalp topographies for HF-to-brain interplay show positive colormaps, indicating that coupling coefficients during emotion elicitation have higher amplitudes than those during the resting state. HF-to-delta (41/60 trials averaged) presented activations in midline frontal and occipitoparietal electrodes, similar to HF-to-theta (46/60 trials averaged). Activations in the HF-to-alpha (45/60 trials averaged) were slightly lateralized to the right hemisphere in frontocentral electrodes and to the left hemisphere in the occipitoparietal region. HF-to-beta (42/60 trials averaged) interplay presents activations in midline frontal and left occipitoparietal region. HF-to-gamma (31/60 trials averaged) presents activations in frontal and occipital regions, with no lateralization. Scalp topographies of brain-to-heart interplay show negative color maps, which means that coupling coefficients during emotion elicitation are lower than in the resting state period, the opposite of ascending modulations. The central electrodes presented a higher absolute amplitude change in all EEG frequency bands: delta-to-HF (42/60 trials averaged), theta-to-HF (45/60 trials averaged), alpha-to-HF (47/60 trials averaged), beta-to-HF (47/60 trials averaged), and gamma-to-HF (41/60 trials averaged). These results indicate that bi-directional modulations between the heart and the brain present significant changes under emotion elicitation in relation to the resting state, with a major change associated with HF oscillations from the heart rate.

**Figure 2.**
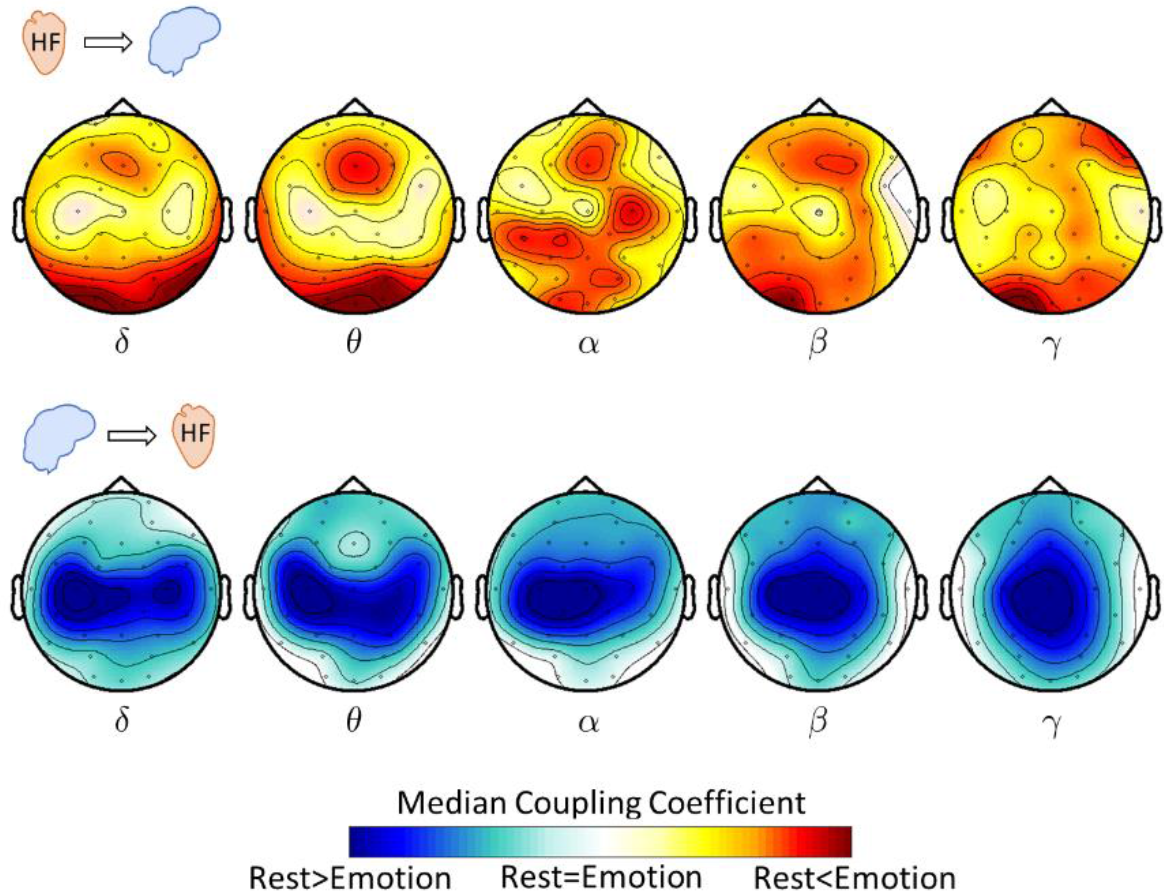
Scalp regions where major changes are observed in HF-to-brain and brain-to-HF interplay during emotion elicitation, in both, the DEAP and MAHNOB datasets. Topographies are constructed by computing the group-median per trial and z-scored among channels before computing the trial-median. Coefficients are previously combined in the time domain by averaging the intervals in which the cluster permutation analysis found a significant change, per trial.

### 2. Correlation analysis between emotions’ self-assessment scores and directional brain-heart coefficients

We explored group-wise correlations of the SDG coefficients with the reported arousal and valence scores. We found in the *DEAP* dataset that delta, theta, and gamma bands receive significant modulations from the HF band, which are correlated to the reported arousal from the self-assessment. As presented in Figure 3A, the HF-to-delta and HF-to-theta bands are related to the arousal in overlapped regions, mainly in the frontal and occipitoparietal electrodes, whereas HF-to-gamma relates to the temporal scalp regions. Figure 3B presents the median SDG coefficients per trial among subjects relating to the median arousal. HF-to-delta interplay was significantly correlated in the electrodes located in the occipitoparietal, left-central, and frontal electrodes (Spearman correlation, R = 0.4802, p = 0.0017). HF-to-theta was correlated with arousal in the occipitoparietal and frontal electrodes (Spearman correlation, R = 0.5205, p = 0.0006) and HF-to-gamma in both temporal regions (Spearman correlation, R = 0.5660, p = 0.0001). As shown in Figure 3C, the high correlation found in the coupling coefficients was not present in the separated components HF (Spearman correlation, R = −0.3039, p = 0.0566), delta band (Spearman correlation, R = 0.0673, p = 0.6800), theta band (Spearman correlation, R = 0.0516, p = 0.7518), and gamma band (Spearman correlation, R = 0.1788, p = 0.2696. In the *DEAP* dataset, we did not find correlations between scores in self-assessment of emotions and SDG coefficients assessing LF-to-brain, brain-to-LF, and brain-to-HF modulations on the arousal and valence dimensions (see the supplementary material for details). We did not find any such relationships in the *MAHNOB* dataset.

**Figure 3.**
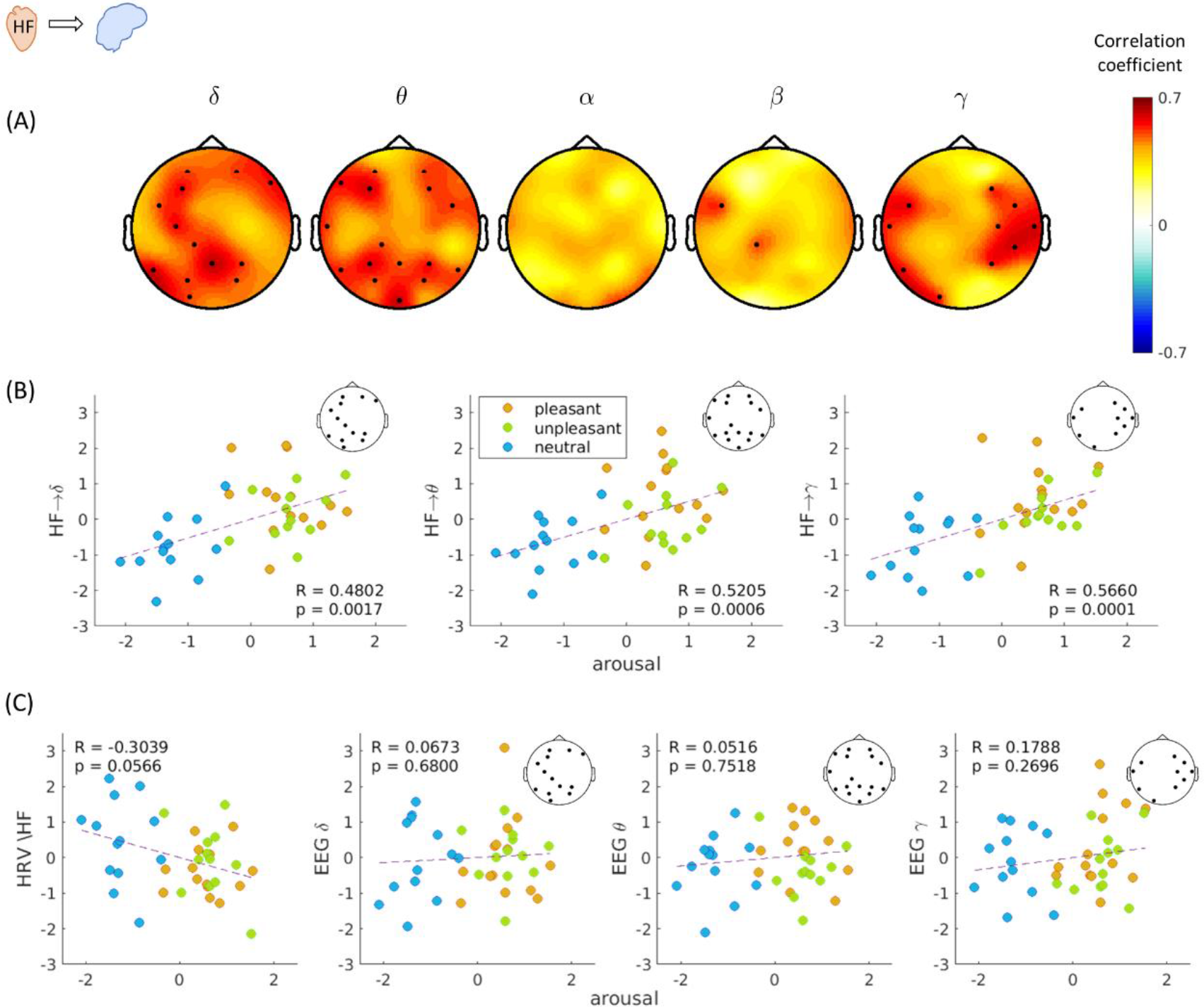
Correlation analysis between ascending SDG coefficients and reported arousal in the self-assessment. Each point evaluated in the correlation analysis corresponds to the group-median value per trial, for reported arousal and SDG coefficients. Individual values were previously normalized within the 40 trials for each subject. (A) Correlation between trial’s median arousal and HF-to-brain median coefficients. Colormaps indicate correlation coefficients (R), and thick electrodes indicate significant correlation corrected for multiple comparisons among channels (p < 0.05/32). (B) Relation between arousal and the coupling coefficient for HF→δ, HF→θ, and HF→γ in the channels where a significant correlation was found. (C) Relation between arousal and HF, EEG δ, EEG θ, and EEG γ.

### 3. Temporal dynamics of the directional brain-heart coefficients under emotion elicitation

We further investigated the overall change in time of the SDG coefficients. For this, we performed group-wise Friedman tests to compare between the resting state and emotion elicitation at different latencies, among all-trials-median. The Friedman test results indicated that significant changes occurred among the conditions presented for both ascending and descending pathways. In Figure 4A, heart-to-brain interplay presents most changes in the HF-to-theta, and in brain-to-heart interplay, these changes are in the alpha-to-HF interplay in the two datasets studied (see the supplementary material for changes in other frequency bands). These results are in line with the outputs of the cluster-based permutation analysis in Figure 2, in which the main changes observed were in HF-to-theta interplay in the frontal and occipitoparietal regions and alpha-to-HF interplay in the central region.

**Figure 4.**
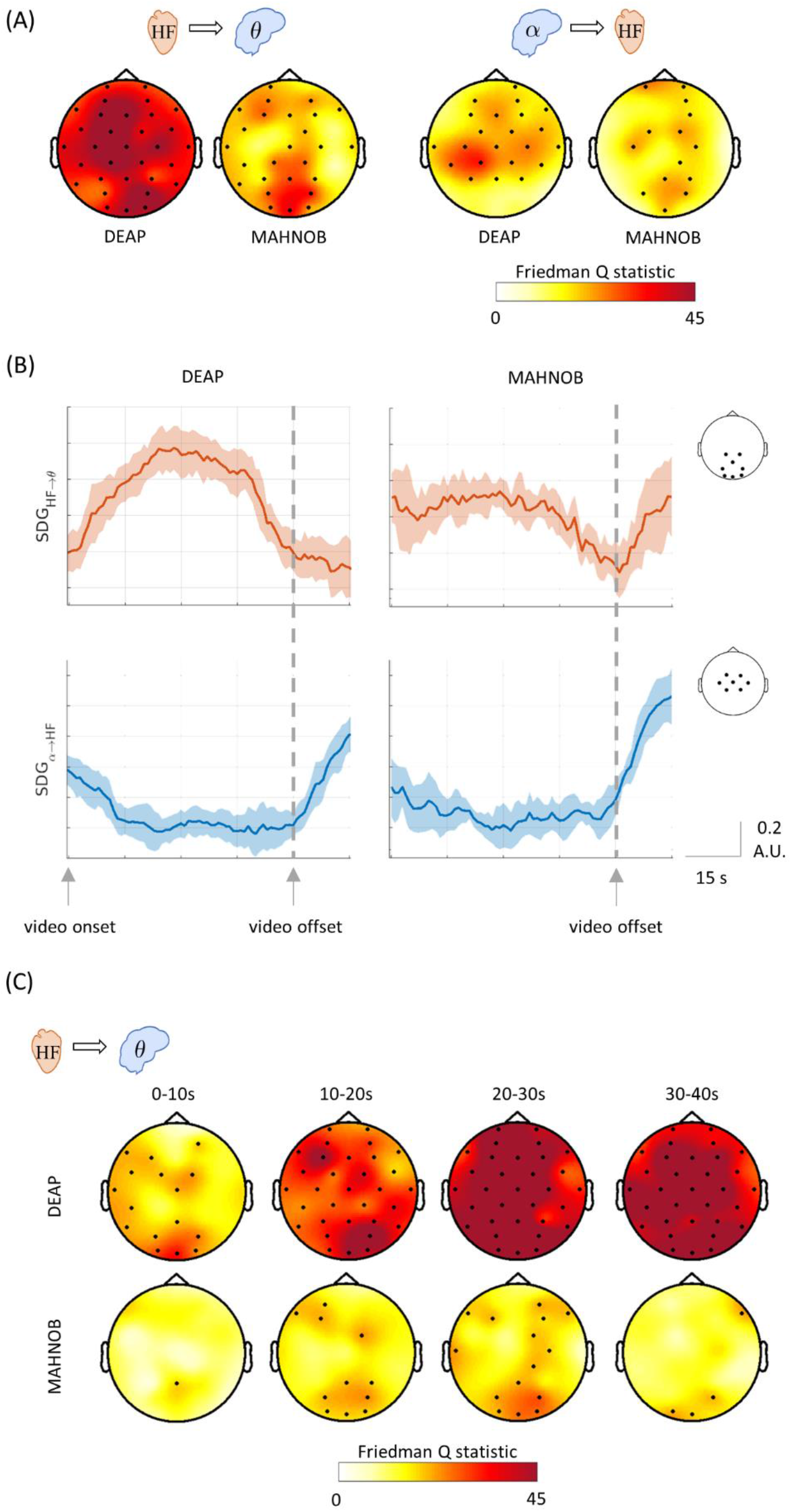
Overall changes in the brain-heart interplay during emotion elicitation. (A) Friedman test of HF-to-theta and alpha-to-HF performed over four averaged time intervals: rest and first three-quarters of the trial, all trials averaged separately for the two datasets. Colormaps indicate the amplitudes of the respective statistics and thick electrodes indicate significance (p < 0.05/32). (B) Averaged time courses among all trials of HF-to-theta and alpha-to-HF coefficients. All trials were aligned at the video offset. Friedman test in four different time windows averaged in four different conditions: resting state and emotion elicitation trials (pleasant, unpleasant, and neutral). Trials of each category were averaged for each subject, separately for the two datasets. Colormaps indicate the amplitudes of the respective statistics and thick electrodes indicate significance (p < 0.05/32).

In Figure 4B, we present the group-median time course aligned in the trial offset. HF-to-theta interplay indicates an overall increase in the SDG coefficient amplitudes during the video visualization, whereas alpha-to-HF in the opposite direction presents a decay toward the video end. The BHI temporal dynamics show that the physiological processes involved in emotion elicitation cause changes in both heart-to-brain and brain-to-heart interplay, with an anticorrelated behavior during the video visualization period. We further investigated the electrodes with major changes in their SDG coefficients using the Friedman test. Figure 4C presents the results for the four different latencies of the video visualization. The four conditions compared in each latency were averaged resting state, pleasant, unpleasant, and neutral trials. The major differences in the ascending pathway appear in the HF-to-theta in regions similar to those observed previously, with a focus on occipitoparietal electrodes in both datasets, being stronger in the *DEAP* dataset. The time window in which the maximum Friedman statistic is found is in the interval 20–30 s with respect to the trial onset in both datasets, indicating that at 20 s, an increase in the variability in the brain-heart coupling occurs for the different types of emotions. The descending pathway from brain-to-HF did not present significant changes according to the Friedman test in the studied time intervals.

Confirming previous findings, Figure 5 presents an overall representation of the *DEAP* dataset, in which the trials’ counts (separated as unpleasant, pleasant, and neutral) presented a significant change in relation to the resting state at different latencies in the cluster-based permutation analysis. It should be noted that it was not possible to draw an analogous figure for the *MANHOB* dataset because different videos had different durations. It can be observed that the 20–30 s latency presents major differences between unpleasant, pleasant, and neutral trials, as previously presented in Figure 4C. We can also observe that these results are in line with the relation to the level of arousal previously presented in Figure 3, as the pleasant and unpleasant emotions present more trials with a significant change overall when compared to the neutral trials. Among all emotion types, heart-to-brain interplay occurs primarily earlier, compared to brain-to-heart interplay. However, brain-to-heart interplay remained longer toward the end of the video visualization. As previously mentioned, the HF power interacts with brain oscillations in both the ascending and descending pathway. However, LF power is also involved in the ascending pathway, as observed mainly in pleasant and unpleasant trials in later stages of video clip visualizations.

**Figure 5.**
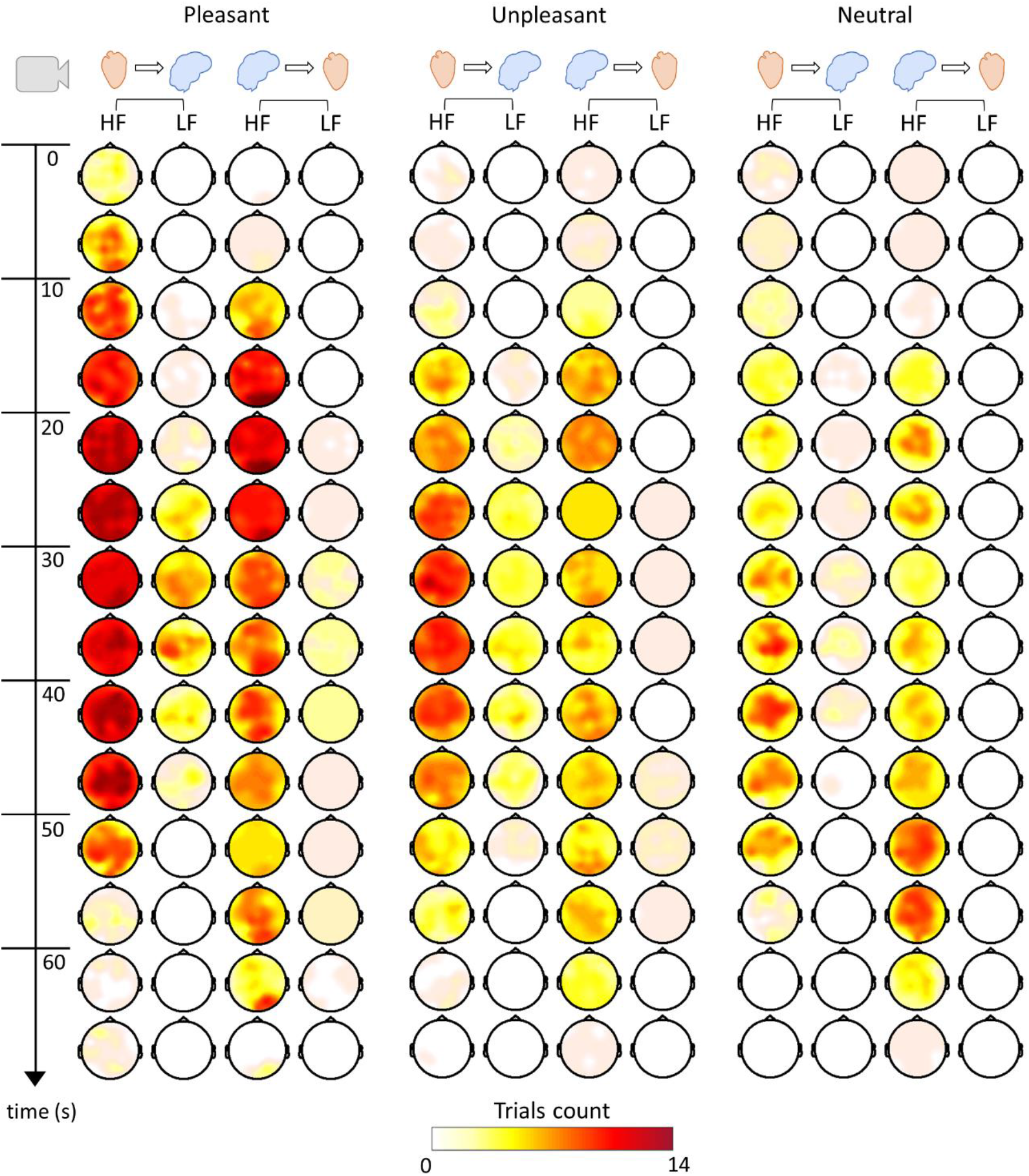
Brain-Heart Interplay time course trial visualization. The scalp topographies displayed correspond to HF-to-brain, LF-to-brain, brain-to-HF, and brain-to-LF. Colormaps indicate the number of videos in which the 5 seconds group-median SDG coefficients during the emotion elicitation have been found significant compared to the rest coupling coefficients previously subtracted per subject.

### 4. Exemplary brain-heart dynamics for pleasant and unpleasant trials

Finally, we present two examples to better illustrate the brain-heart dynamics contrasting a pleasant and unpleasant emotion. To do so, we present in Figure 6 the evolution of SDG coefficients at different latencies (to visualize an exemplary case for the SDG time courses for one subject, see the supplementary material). In Figure 6A, we observed an example taken from the *DEAP* dataset corresponding to an extract of the music video “The One I Once Was,” by the Norwegian band *Mortemia*. We placed this music video in the group of unpleasant trials based on the valence and arousal self-assessment performed by the participants in this study. The video starts with a close-up of an executioner where activation is immediately observed on the parietal scalp region in the HF-to-brain interplay, followed by an overall scalp change from the brain-to-HF interplay. The video continues with the execution of a man, in which the whole process causes an overall activation of the scalp in the HF-to-brain interplay, still focused on parietal electrodes. Around the 30^th^second, the man is executed, and a higher coupling appears on frontal electrodes in the HF-to-brain interplay, an overall scalp activation of lower amplitude appears in the LF-to-brain interplay, and the brain-to-HF interplay is maintained with a stronger coupling on central electrodes. The video is followed by a deceased man accompanied by two nuns and a cemetery scene, where a decrease in the coupling coefficients is observed with the HF-to-brain, which lasts longer.

**Figure 6.**
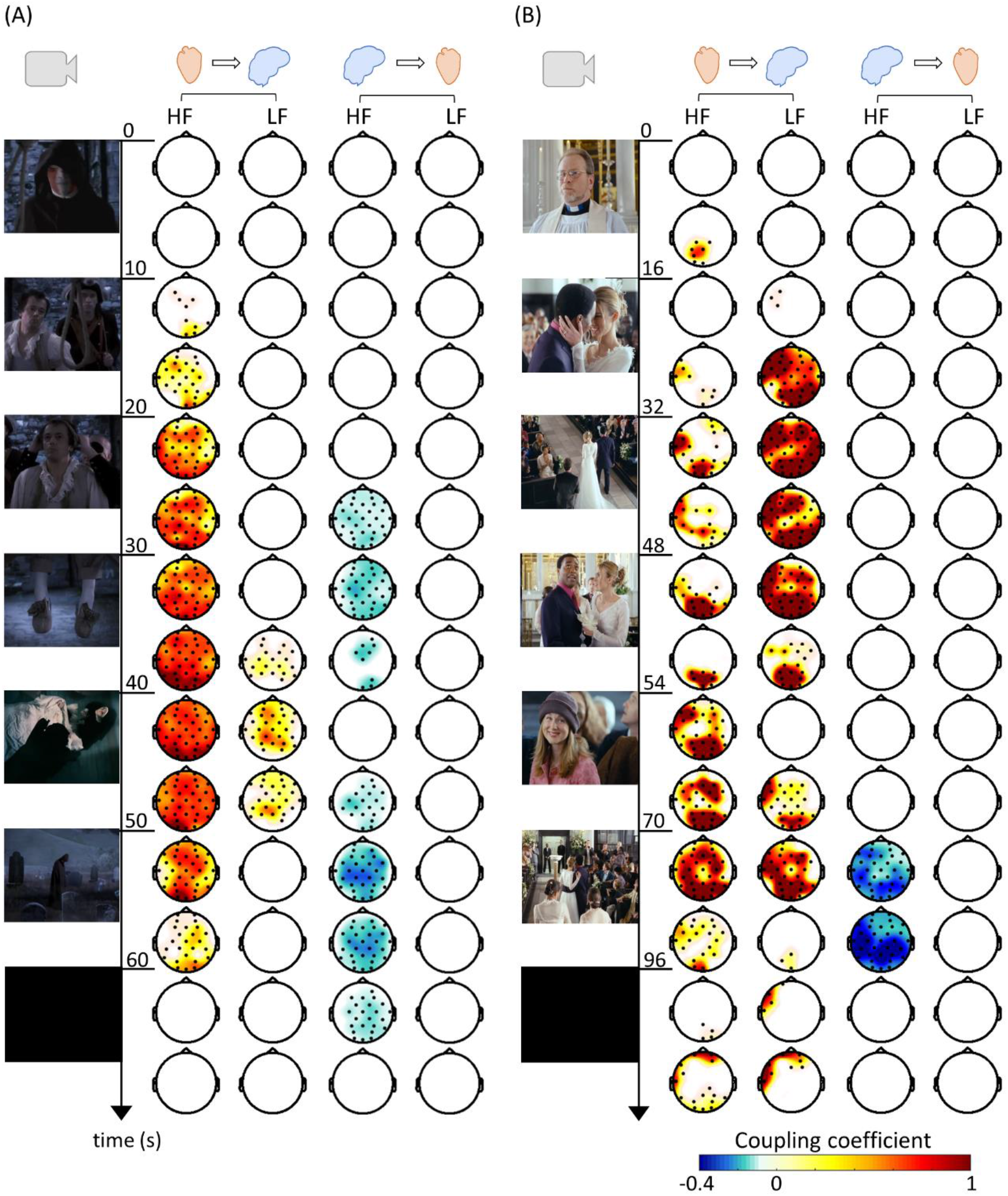
Brain-Heart Interplay time course trial visualization. The scalp topographies displayed correspond to HF-to-brain, LF-to-brain, brain-to-HF, and brain-to-LF. Colormaps indicate the 5 seconds group-median coupling coefficients during the emotion elicitation, with the rest coupling coefficients previously subtracted per subject. A positive coupling coefficient means that the emotion > rest, and a negative coupling coefficient means emotion < rest. The coupling coefficients were set to zero in the samples where non-significant clusters are found. The information per channel within the 5 seconds was wrapped taking the maximum absolute value among the five frequency bands. (A) Trial 31 from DEAP dataset. Screen captures from “The One I Once Was” music video by Mortemia, Napalm Records. (B) Trial 9 from the MAHNOB dataset. Screen captures from the 2003 film “Love Actually,” by Richard Curtis, Universal Pictures.

In Figure 6B, we observed an example taken from the *MAHNOB* dataset corresponding to an extract of the 2003 film “Love Actually,” a romantic comedy by *Richard Curtis*. We placed this trial in the group of pleasant trials based on valence and arousal self-assessments performed by the participants of this study. The video starts with a priest finishing a wedding ceremony; when the applause and instrumental wedding music begin, the man kisses the bride, and activation on the parietal region in the HF-to-brain interplay is observed. LF-to-brain interplay appears in the left frontal scalp region when a short dialog occurs, followed by the start of choral music and progressively appearing musicians from random positions inside the church, playing a cover version of “All You Need Is Love,” by *The Beatles*. During the music, strong activations in the frontal and parietal electrodes are observed in the HF-to-brain interplay, finishing with a guitar solo in which a brain-to-HF interplay is observed as well.

## Discussion and Conclusion

The ANS activity, comprising the heart rate and other visceral activities, has been widely associated with emotional processing,^35^ leading to a century-long debate on the role of the ANS in emotions.^1^Recent studies have uncovered that ascending inputs from the heart are involved in essential aspects of cognition, such as subjective perception, self-awareness, and consciousness.^7^We used a mathematical modeling approach^25^ to represent the interplay between the physiological components gathered from the brain and the heart during emotion elicitation. The results show that cardiac parasympathetic activity plays a causal role in emotional processing, sustaining both ascending and descending brain-heart interactions. We found that ascending BHI measurements are correlated to perceived arousal, and concurrent bi-directional communication between the brain and body sustains emotional processing through specific timings.

We observed that HF power, a marker of parasympathetic activity, is actively involved in bidirectional BHI during emotion elicitation. The variation of the parasympathetic tone has been described before to be related to processes of fluctuations in attention and emotional processing.^36^These autonomic markers have shown their capacity to describe emotions and related states in healthy participants^35^as well as under pathological conditions.^37^ Autonomic outflow has been previously related to behavior in polyvagal theory,^38^ describing neural circuits involved in homeostatic regulation and adaptation. Polyvagal theory describes the interaction of sympathetic and parasympathetic nervous systems, in which the parasympathetic branch is associated with emotion regulation because of its behavioral correlates with reactivity, the expression of emotion, and self-regulation skills.^38^The mediating role of the vagus nerve during emotional processing fits with the communication loops observed in the brain-heart interplay in our results. However, beyond mediation, we observed that parasympathetic activity initiates emotional processing. The observed ascending pathway of vagal activity towards the brain corroborates previous findings that heartbeats shape brain dynamics during subjective perception.^4,39^Therefore, this could imply emotions are not the result of an interpretation of peripheral physiological changes but rather an integration of these inputs in the brain, from the visual, auditory, and somatosensory perception stages to the ongoing evolution of subjective experiences. These results support the hypothesis that the heart contributes to a first-person point of view for conscious/subjective experiences.^7^

The causal nature of ANS activity in emotions is consistent with physiological modeling and experimental data, as neurovisceral integration models indicate that neural circuits integrate central and autonomic emotional responses, with dynamic contextual adaptations.^8,14,40^In theoretical developments, the specific role of bodily signals in subjectivity and consciousness has been conceived in different ways from the causation viewpoint. The somatic marker hypothesis of *Damasio*^41^affirms that the meta-representation of bodily states constitutes an emotional feeling, generating a gut feeling, which influences cognitive processes. For some authors, visceral activity might provide information for the foundation of emotions.^5,10^ For others, visceral activity is considered a central factor, where the neural monitoring of ascending visceral inputs is inherent for subjective perception, emotions, and consciousness.^7,42^

We showed that vagal modulations to the EEG theta activity primarily occurred during emotion elicitation, and these modulations were correlated with the reported arousal. The group-wise arousal was related to delta and theta bands in the frontal and occipitoparietal scalp regions as well as to the gamma band in temporal areas. To verify whether the results were related exclusively to the BHI, we performed the same analysis separately for brain and heart components, confirming that arousal changes cannot be explained by changes in cardiac parameters or EEG power by themselves. The interactions between EEG activity and heart rate have been described previously in machine learning studies, showing that the combination of these features in classifiers provides additional information related to consciousness and emotions.^43,44^Consistent with previous studies, BHI can describe certain cognitive states, which cannot be explained by changes in cardiac parameters by themselves.^15,21,45,46^In particular, the relationship between heartbeat dynamics and theta band was previously described in the resting state, showing that heartbeats may induce theta synchronizations in defined brain networks.^47^In this study, we showed that the theta band is actively modulated by vagal inputs under emotion elicitation. ^48^

Theta activation has been repeatedly reported in different paradigms, such as the visualization of emotional faces^49^and emotional content in films.^50^Our results corroborate a previous report,^51^in which the midline frontal theta is associated with states of high arousal and high valence, followed by synchronization with posterior scalp areas. Previous studies have described that changes in theta are related to the level of arousal perceived,^50,52^although our results did not show a relationship in the theta band *per se* but rather in its afferent modulation from HF oscillations. The other EEG bands related to arousal were delta and gamma; however, few studies have reported such relations. The delta band has been shown to have the potential to describe emotional processing,^53^and the close relationship between the theta and delta bands during emotional processing has been previously described as well.^54^ With respect to the gamma band, we found that the arousal relation with the ascending modulations in temporal lobes could be somehow related to previous evidence linking theta and gamma bands, as an arousal-dependent synchronization.^49^

Descending modulations were suppressed during the emotion elicitation tasks, particularly in the HF oscillations. The observed modulation involved a wide part of the EEG spectrum, from the alpha to the gamma band, with the alpha range being the most influential. The variations in brain-to-heart coefficients were stronger in the central scalp regions, with a slight extension towards the parietal and temporal areas. In contrast to the relation of alpha oscillations in emotions reported in other studies where it is related mainly to changes in frontal and prefrontal lobe activity.^55^Less evidence is reported about beta and gamma oscillations in emotions; however, these higher frequencies are reported as a synchronized or desynchronized activity with respect to alpha^56^or theta bands.^49^We could not distinguish the different types of emotions using information from the descending pathway, as we did not find associations with arousal or valence. However, we found that descending modulations were decreased in most of the trials compared to the resting state. Furthermore, descending changes were more delayed compared to the ascending pathway. Here, we hypothesized the possible involvement of these physiological changes. There is an integral relationship between emotions and other cognitive processes, which may imply that emotions can shape other cognitive processes, such as attention, memory, and decision making.^57,58^ Previous experimental results support this claim, for example, the late or delayed changes observed in the EEG oscillations, suggesting post-processing of information, such as EEG beta band in attention tasks,^59^or delayed power variations of the gamma band during emotional elicitations.^60,61^ Some authors discuss that emotional experiences involve a posterior process of re-experiencing, in which perceptual, somatovisceral, and motor mechanisms may be involved.^62,63^The functional activity of alpha, beta, and gamma bands in the central and parietal regions have been previously described in motor and somatosensory activity.^64–66^ Studies on the somatosensory experience of emotions state that bodily sensations are felt in different body parts as a function of the type of emotions felt.^67^In fact, the experience of others’ emotions may be closely related to the activity in the somatosensory cortex as well.^68^The specific role of the descending pathway remains to be further investigated to determine whether it relates to bodily re-experiencing, or to aspects other than the emotion itself but rather more general aspects, such as perception, attention, or memory.^57,58^

Several studies focused on BHI and emotions in different paradigms have been conducted. To date, available evidence on BHI and emotions describes dynamic and non-linear interactions.^8,22–24^ Further evidence exists in studies on heartbeat-evoked potentials; nevertheless, there is a lack of convergence in these results.^21^For instance, heartbeat-evoked potentials were correlated with alpha power during the task with audio-visual stimuli.^69^However, as mentioned above, our results show that the SDG model in ascending modulations relates to the specific perceived arousal, and alpha activity is more related to the descending pathway.

Other studies have reported heartbeat-evoked potentials to valence.^70,71^ As these interoceptive markers show some variability in the different paradigms, it remains unclear whether BHI measured from heartbeat-evoked potentials actually describes aspects of valence, arousal, or other subjective aspects involved in emotion, such as ownership, motivation, dominance, or abnormalities in emotional processing.^21^ However, evidence in pathological conditions showed a relationship between disrupted emotion recognition and heartbeat-evoked potentials, indicating altered interoceptive mechanisms.^39^ We repeatedly found two well-defined scalp regions in the midline frontal and occipitoparietal electrodes, in which ascending SDG coefficients were concisely related to emotion elicitation. These results suggest the possibility of multiple brain nodes participating in the ongoing process. Neuroimaging studies have shown evidence of the DMN and the processing of self-relevant and affective decisions,^16^with the main nodes in the medial prefrontal cortex and posterior cingulate cortex. The fact that BHI is related to the DMN, in aspects including autonomic regulation^14^and interoception in self-related cognition,^15^suggests that it may also be involved in emotions. The DMN, which is usually associated with passive states,^13^presents an activation/deactivation behavior that seems to be related to the switch between inward mental activity and outward goal-directed activity.^72^How the switching activity occurs is yet to be completely understood, but increasing evidence supports that the monitoring of the peripheral neural activity by the DMN may turn the brain to a goal and externally oriented mode in specific circumstances.^72^For instance, the functions of the different nodes of the DMN may dissociate depending on which aspect of the self is involved in spontaneous thoughts.^15^In this regard, having DMN node activations as a function of the level of arousal may imply that the higher the arousal, the higher the system is directed from inward out. It is also interesting to note that the CAN and DMN share a part of the brain structures, for example, the medial prefrontal cortex. As a partially alternative framework of interpretation, it is interesting to note that the DMN, usually described as an intrinsic system, has been described alternatively as a network for integration and online updating of experiences.^73^Indeed, bridging the internal state with external relevant information is expected to occur during the emotional experience.

The interplay between the CNS and ANS involves not only the brain and heart but also other bodily signals, such as electrodermal activity, breathing, gastrointestinal activity, or pupil diameter.^74^ We believe that the proposed analysis can be enriched in future works at a multisystem level. Furthermore, even though we employed two large and publicly available datasets,^26,27^one may argue about how our results are bound to the specific emotional elicitation (video presentation) and how BHI might vary according to different emotional elicitation techniques, which represent an interesting further line of investigation. Although the SDG model cannot assess spontaneous neural responses in a millisecond resolution, as heartbeat-evoked potentials do, it has the advantage of assessing ongoing, continuous, and bi-directional modulations.

The tight relationship between well-being and autonomic function is evidenced in cases of dysfunctional brain-heart interplay in psychopathological conditions.^75^Promising developments may lead to a better understanding of emotional processing and bodily states, as clinical applications involving the analysis of brain-heart interactions have brought overall important advances.^44,76,77^Beyond mental and neurological health, a vast amount of evidence shows that other bodily activities actively react as a function of ongoing brain activity, including gut activity, endocrine responses, and inflammation,^6^ indicating the importance of neural homeostasis in the human body. Our results may be expanded to better clarify the role of the BHI in the mutual vulnerability between mental and physical conditions, and thus provide a psychophysiological model of how physical health may contribute to mental health risk factors, and vice versa.

We have shown how the dynamic interplay between the central and autonomic peripheral nervous systems sustains emotional experiences through specific timings and cortical areas. During emotional elicitation, we have shown how autonomic dynamics on cardiovascular control initiate the physiological response to emotion in the direction of the CNS, possibly belonging to the DMN and CAN. This activity is correlated with the subjective experience of arousal. Our results add new momentum to the theory of emotions, suggesting that peripheral neural dynamics of the cardiovascular ANS may trigger the experience of emotions at the brain level.

## References

1. Pace-Schott, E. F. et al. Physiological feelings. Neuroscience & Biobehavioral Reviews 103, 267–304 (2019).

2. Chen, W. G. et al. The Emerging Science of Interoception: Sensing, Integrating, Interpreting, and Regulating Signals within the Self. Trends in Neurosciences 44, 3–16 (2021).

3. Damasio, A. R. et al. Subcortical and cortical brain activity during the feeling of self-generated emotions. Nature Neuroscience 3, 1049–1056 (2000).

4. Park, H.-D., Correia, S., Ducorps, A. & Tallon-Baudry, C. Spontaneous fluctuations in neural responses to heartbeats predict visual detection. Nat. Neurosci. 17, 612–618 (2014).

5. Craig, A. D. How do you feel — now? The anterior insula and human awareness. Nature Reviews Neuroscience 10, 59–70 (2009).

6. Critchley, H. D. & Garfinkel, S. N. Interoception and emotion. Current Opinion in Psychology 17, 7–14 (2017).

7. Azzalini, D., Rebollo, I. & Tallon-Baudry, C. Visceral Signals Shape Brain Dynamics and Cognition. Trends in Cognitive Sciences 23, 488–509 (2019).

8. Thayer, J. F. & Lane, R. D. A model of neurovisceral integration in emotion regulation and dysregulation. J Affect Disord 61, 201–216 (2000).

9. Cameron, O. G. Interoception: The Inside Story—A Model for Psychosomatic Processes. Psychosomatic Medicine 63, 697–710 (2001).

10. Craig, A. D. How do you feel? Interoception: the sense of the physiological condition of the body. Nat. Rev. Neurosci. 3, 655–666 (2002).

11. Beissner, F., Meissner, K., Bär, K.-J. & Napadow, V. The autonomic brain: an activation likelihood estimation meta-analysis for central processing of autonomic function. J. Neurosci. 33, 10503–10511 (2013).

12. Valenza, G. et al. The central autonomic network at rest: Uncovering functional MRI correlates of time-varying autonomic outflow. NeuroImage 197, 383–390 (2019).

13. Raichle, M. E. The Brain’s Default Mode Network. Annual Review of Neuroscience 38, 433–447 (2015).

14. Thayer, J. F., Ahs, F., Fredrikson, M., Sollers, J. J. & Wager, T. D. A meta-analysis of heart rate variability and neuroimaging studies: implications for heart rate variability as a marker of stress and health. Neurosci Biobehav Rev 36, 747–756 (2012).

15. Babo-Rebelo, M., Richter, C. G. & Tallon-Baudry, C. Neural Responses to Heartbeats in the Default Network Encode the Self in Spontaneous Thoughts. J. Neurosci. 36, 7829–7840 (2016).

16. Andrews-Hanna, J. R., Reidler, J. S., Sepulcre, J., Poulin, R. & Buckner, R. L. Functional-anatomic fractionation of the brain’s default network. Neuron 65, 550–562 (2010).

17. Critchley, H. D., Corfield, D. R., Chandler, M. P., Mathias, C. J. & Dolan, R. J. Cerebral correlates of autonomic cardiovascular arousal: a functional neuroimaging investigation in humans. J Physiol 523, 259–270 (2000).

18. Phan, K. L., Wager, T., Taylor, S. F. & Liberzon, I. Functional neuroanatomy of emotion: a meta-analysis of emotion activation studies in PET and fMRI. Neuroimage 16, 331–348 (2002).

19. Anders, S. et al. Parietal somatosensory association cortex mediates affective blindsight. Nature Neuroscience 7, 339–340 (2004).

20. Candia-Rivera, D., Catrambone, V. & Valenza, G. The Role of Electroencephalography Electrical Reference in the Assessment of Functional Brain–Heart Interplay: From Methodology to User Guidelines. Journal of Neuroscience Methods Under review, (2021).

21. Park, H.-D. & Blanke, O. Heartbeat-evoked cortical responses: Underlying mechanisms, functional roles, and methodological considerations. NeuroImage 197, 502–511 (2019).

22. Catrambone, V., Greco, A., Scilingo, E. P. & Valenza, G. Functional Linear and Nonlinear Brain–Heart Interplay during Emotional Video Elicitation: A Maximum Information Coefficient Study. Entropy 21, 892 (2019).

23. Greco, A. et al. Lateralization of directional brain-heart information transfer during visual emotional elicitation. American Journal of Physiology-Regulatory, Integrative and Comparative Physiology 317, R25–R38 (2019).

24. Lewis, M. D. Bridging emotion theory and neurobiology through dynamic systems modeling. Behavioral and Brain Sciences 28, 169–194 (2005).

25. Catrambone, V., Greco, A., Vanello, N., Scilingo, E. P. & Valenza, G. Time-Resolved Directional Brain-Heart Interplay Measurement Through Synthetic Data Generation Models. Ann Biomed Eng 47, 1479–1489 (2019).

26. Koelstra, S. et al. DEAP: A Database for Emotion Analysis ;Using Physiological Signals. IEEE Transactions on Affective Computing 3, 18–31 (2012).

27. Soleymani, M., Lichtenauer, J., Pun, T. & Pantic, M. A Multimodal Database for Affect Recognition and Implicit Tagging. IEEE Transactions on Affective Computing 3, 42–55 (2012).

28. Oostenveld, R., Fries, P., Maris, E. & Schoffelen, J.-M. FieldTrip: Open Source Software for Advanced Analysis of MEG, EEG, and Invasive Electrophysiological Data. Computational Intelligence and Neuroscience 2011, 9 pages (2011).

29. Gabard-Durnam, L. J., Mendez Leal, A. S., Wilkinson, C. L. & Levin, A. R. The Harvard Automated Processing Pipeline for Electroencephalography (HAPPE): Standardized Processing Software for Developmental and High-Artifact Data. Front. Neurosci. 12, (2018).

30. Citi, L., Brown, E. N. & Barbieri, R. A Real-Time Automated Point Process Method for Detection and Correction of Erroneous and Ectopic Heartbeats. IEEE Trans Biomed Eng 59, 2828–2837 (2012).

31. Orini, M., Bailón, R., Mainardi, L. T., Laguna, P. & Flandrin, P. Characterization of dynamic interactions between cardiovascular signals by time-frequency coherence. IEEE Trans Biomed Eng 59, 663–673 (2012).

32. Brennan, M., Palaniswami, M. & Kamen, P. Poincaré plot interpretation using a physiological model of HRV based on a network of oscillators. Am. J. Physiol. Heart Circ. Physiol. 283, H1873-1886 (2002).

33. Al-Nashash, H., Al-Assaf, Y., Paul, J. & Thakor, N. EEG signal modeling using adaptive Markov process amplitude. IEEE Transactions on Biomedical Engineering 51, 744–751 (2004).

34. Maris, E. & Oostenveld, R. Nonparametric statistical testing of EEG- and MEG-data. Journal of Neuroscience Methods 164, 177–190 (2007).

35. Kreibig, S. D. Autonomic nervous system activity in emotion: A review. Biological Psychology 84, 394–421 (2010).

36. Thayer, J. F. & Siegle, G. J. Neurovisceral integration in cardiac and emotional regulation. IEEE Engineering in Medicine and Biology Magazine 21, 24–29 (2002).

37. Vazquez, L. et al. High frequency heart-rate variability predicts adolescent depressive symptoms, particularly anhedonia, across one year. J Affect Disord 196, 243–247 (2016).

38. Porges, S. W. The polyvagal perspective. Biol Psychol 74, 116–143 (2007).

39. Salamone, P. C. et al. Interoception primes emotional processing: Multimodal evidence from neurodegeneration. J. Neurosci. (2021) doi:10.1523/JNEUROSCI.2578-20.2021.

40. Hagemann, D., Waldstein, S. R. & Thayer, J. F. Central and autonomic nervous system integration in emotion. Brain and Cognition 52, 79–87 (2003).

41. Damasio, A. The feeling of what happens: Body and emotion in the making of consciousness. xiv, 386 (Harcourt College Publishers, 1999).

42. Smith, R., Thayer, J. F., Khalsa, S. S. & Lane, R. D. The hierarchical basis of neurovisceral integration. Neuroscience & Biobehavioral Reviews 75, 274–296 (2017).

43. Marín-Morales, J. et al. Affective computing in virtual reality: emotion recognition from brain and heartbeat dynamics using wearable sensors. Scientific Reports 8, 13657 (2018).

44. Candia-Rivera, D. et al. Neural responses to heartbeats detect residual signs of consciousness during resting state in post-comatose patients. J. Neurosci. (2021) doi:10.1523/JNEUROSCI.1740-20.2021.

45. Naji, M., Krishnan, G. P., McDevitt, E. A., Bazhenov, M. & Mednick, S. C. Coupling of autonomic and central events during sleep benefits declarative memory consolidation. Neurobiol Learn Mem 157, 139–150 (2019).

46. Chen, P.-C., Whitehurst, L. N., Naji, M. & Mednick, S. C. Autonomic/central coupling benefits working memory in healthy young adults. Neurobiology of Learning and Memory 173, 107267 (2020).

47. Kim, J. & Jeong, B. Heartbeat Induces a Cortical Theta-Synchronized Network in the Resting State. eNeuro 6, ENEURO.0200-19.2019 (2019).

48. Patron, E., Mennella, R., Messerotti Benvenuti, S. & Thayer, J. F. The frontal cortex is a heart-brake: Reduction in delta oscillations is associated with heart rate deceleration. NeuroImage 188, 403–410 (2019).

49. Balconi, M. & Pozzoli, U. Arousal effect on emotional face comprehension: Frequency band changes in different time intervals. Physiology & Behavior 97, 455–462 (2009).

50. Krause, C. M., Viemerö, V., Rosenqvist, A., Sillanmäki, L. & Aström, T. Relative electroencephalographic desynchronization and synchronization in humans to emotional film content: an analysis of the 4-6, 6-8, 8-10 and 10-12 Hz frequency bands. Neurosci Lett 286, 9–12 (2000).

51. Aftanas, L. I. & Golocheikine, S. A. Human anterior and frontal midline theta and lower alpha reflect emotionally positive state and internalized attention: high-resolution EEG investigation of meditation. Neurosci Lett 310, 57–60 (2001).

52. Aftanas, L. I., Varlamov, A. A., Pavlov, S. V., Makhnev, V. P. & Reva, N. V. Time-dependent cortical asymmetries induced by emotional arousal: EEG analysis of event-related synchronization and desynchronization in individually defined frequency bands. International Journal of Psychophysiology 44, 67–82 (2002).

53. Klados, M. A. et al. A framework combining delta Event-Related Oscillations (EROs) and Synchronisation Effects (ERD/ERS) to study emotional processing. Comput Intell Neurosci 549419 (2009) doi:10.1155/2009/549419.

54. Knyazev, G. G., Slobodskoj-Plusnin, J. Y. & Bocharov, A. V. Event-related delta and theta synchronization during explicit and implicit emotion processing. Neuroscience 164, 1588–1600 (2009).

55. Allen, J. J. B., Coan, J. A. & Nazarian, M. Issues and assumptions on the road from raw signals to metrics of frontal EEG asymmetry in emotion. Biological Psychology 67, 183–218 (2004).

56. Jessen, S. & Kotz, S. A. The temporal dynamics of processing emotions from vocal, facial, and bodily expressions. NeuroImage 58, 665–674 (2011).

57. Pessoa, L. On the relationship between emotion and cognition. Nature Reviews Neuroscience 9, 148–158 (2008).

58. LeBlanc, V. R., McConnell, M. M. & Monteiro, S. D. Predictable chaos: a review of the effects of emotions on attention, memory and decision making. Adv Health Sci Educ Theory Pract 20, 265–282 (2015).

59. Vázquez Marrufo, M., Vaquero, E., Cardoso, M. J. & Gómez, C. M. Temporal evolution of α and β bands during visual spatial attention. Cognitive Brain Research 12, 315–320 (2001).

60. Keil, A. et al. Effects of emotional arousal in the cerebral hemispheres: a study of oscillatory brain activity and event-related potentials. Clinical Neurophysiology 112, 2057–2068 (2001).

61. Aftanas, L. I., Reva, N. V., Varlamov, A. A., Pavlov, S. V. & Makhnev, V. P. Analysis of Evoked EEG Synchronization and Desynchronization in Conditions of Emotional Activation in Humans: Temporal and Topographic Characteristics. Neuroscience and Behavioral Physiology 34, 859–867 (2004).

62. Niedenthal, P. M. Embodying Emotion. Science 316, 1002–1005 (2007).

63. Dum, R. P., Levinthal, D. J. & Strick, P. L. Motor, cognitive, and affective areas of the cerebral cortex influence the adrenal medulla. Proc Natl Acad Sci U S A 113, 9922–9927 (2016).

64. Babiloni, C. et al. Anticipation of somatosensory and motor events increases centro-parietal functional coupling: An EEG coherence study. Clinical Neurophysiology 117, 1000–1008 (2006).

65. Ritter, P., Moosmann, M. & Villringer, A. Rolandic alpha and beta EEG rhythms’ strengths are inversely related to fMRI-BOLD signal in primary somatosensory and motor cortex. Human Brain Mapping 30, 1168–1187 (2009).

66. Nakagawa, K. et al. Neuromagnetic beta oscillation changes during motor imagery and motor execution of skilled movements. NeuroReport 22, 217–222 (2011).

67. Nummenmaa, L., Hari, R., Hietanen, J. K. & Glerean, E. Maps of subjective feelings. PNAS 115, 9198–9203 (2018).

68. Sel, A., Calvo-Merino, B., Tsakiris, M. & Forster, B. The somatotopy of observed emotions. Cortex 129, 11–22 (2020).

69. Luft, C. D. B. & Bhattacharya, J. Aroused with heart: Modulation of heartbeat evoked potential by arousal induction and its oscillatory correlates. Scientific Reports 5, 15717 (2015).

70. Couto, B. et al. Heart evoked potential triggers brain responses to natural affective scenes: A preliminary study. Autonomic Neuroscience 193, 132–137 (2015).

71. Kim, J. et al. Sad faces increase the heartbeat-associated interoceptive information flow within the salience network: a MEG study. Scientific Reports 9, 430 (2019).

72. Riemer, F. et al. Dynamic switching between intrinsic and extrinsic mode networks as demands change from passive to active processing. Scientific Reports 10, 21463 (2020).

73. Yeshurun, Y., Nguyen, M. & Hasson, U. The default mode network: where the idiosyncratic self meets the shared social world. Nature Reviews Neuroscience 22, 181–192 (2021).

74. Rebollo, I., Devauchelle, A.-D., Béranger, B. & Tallon-Baudry, C. Stomach-brain synchrony reveals a novel, delayed-connectivity resting-state network in humans. eLife 7, e33321 (2018).

75. Beauchaine, T. P. & Thayer, J. F. Heart rate variability as a transdiagnostic biomarker of psychopathology. Int J Psychophysiol 98, 338–350 (2015).

76. Terhaar, J., Viola, F. C., Bär, K.-J. & Debener, S. Heartbeat evoked potentials mirror altered body perception in depressed patients. Clin Neurophysiol 123, 1950–1957 (2012).

77. Perogamvros, L. et al. Increased heartbeat-evoked potential during REM sleep in nightmare disorder. NeuroImage: Clinical 22, 101701 (2019).

