## supplementary material for "Cardiac sympathovagal activity initiates a functional brain-body response to emotional processing"

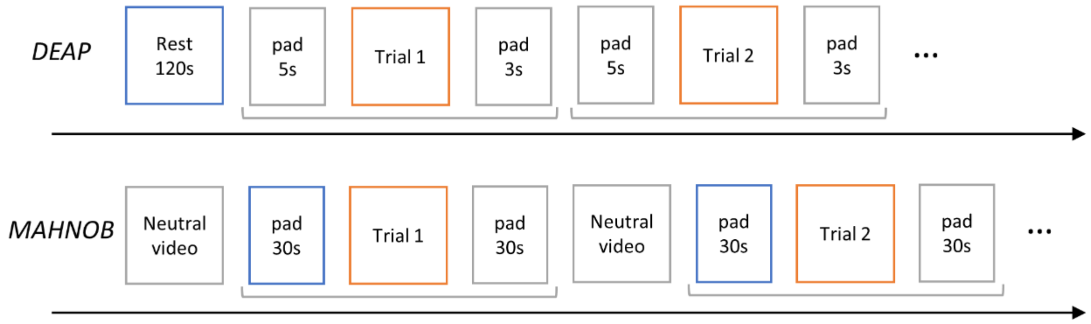


*Figure 1. Simplified protocol description for DEAP and MAHNOB datasets. Red squares indicate the periods when the videos of emotion elicitation are visualized, and the blue squares indicate the rest period used in this study to compare the video trials.*

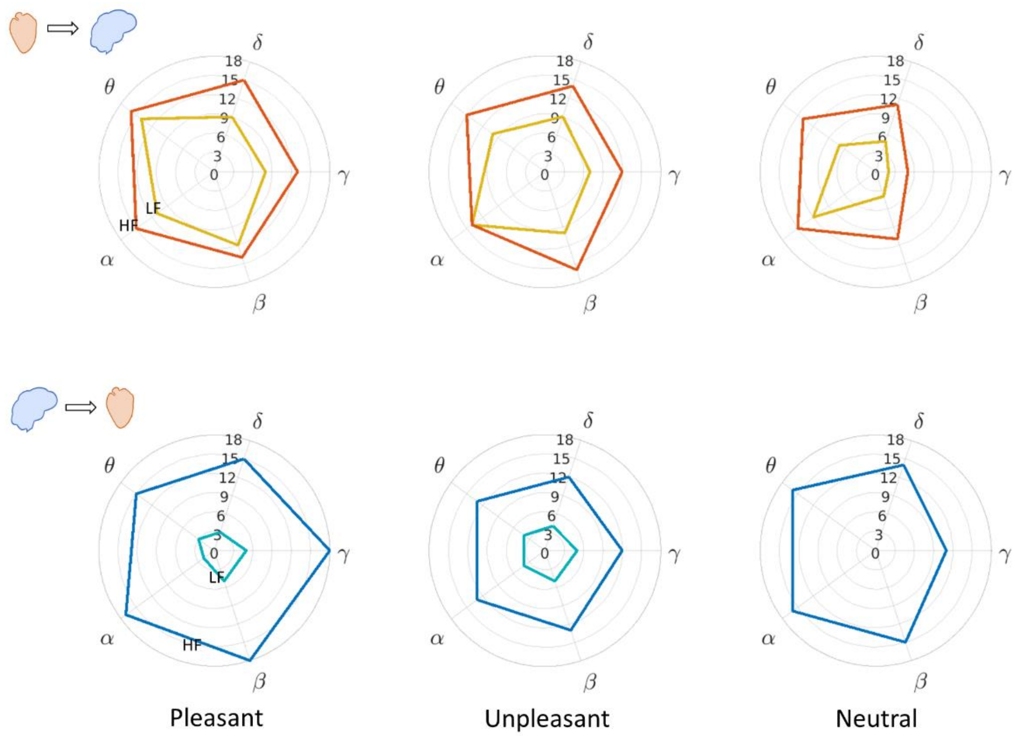


*Figure 2. Relevance of the EEG frequency bands in the Brain-Heart Interplay. The polar histograms indicate the number of trials among the two datasets, for (A) Heart-to-Brain and (B) Brain-to-Heart, in which the respective bands presented a significant change from rest, based on the cluster-based permutation analysis*

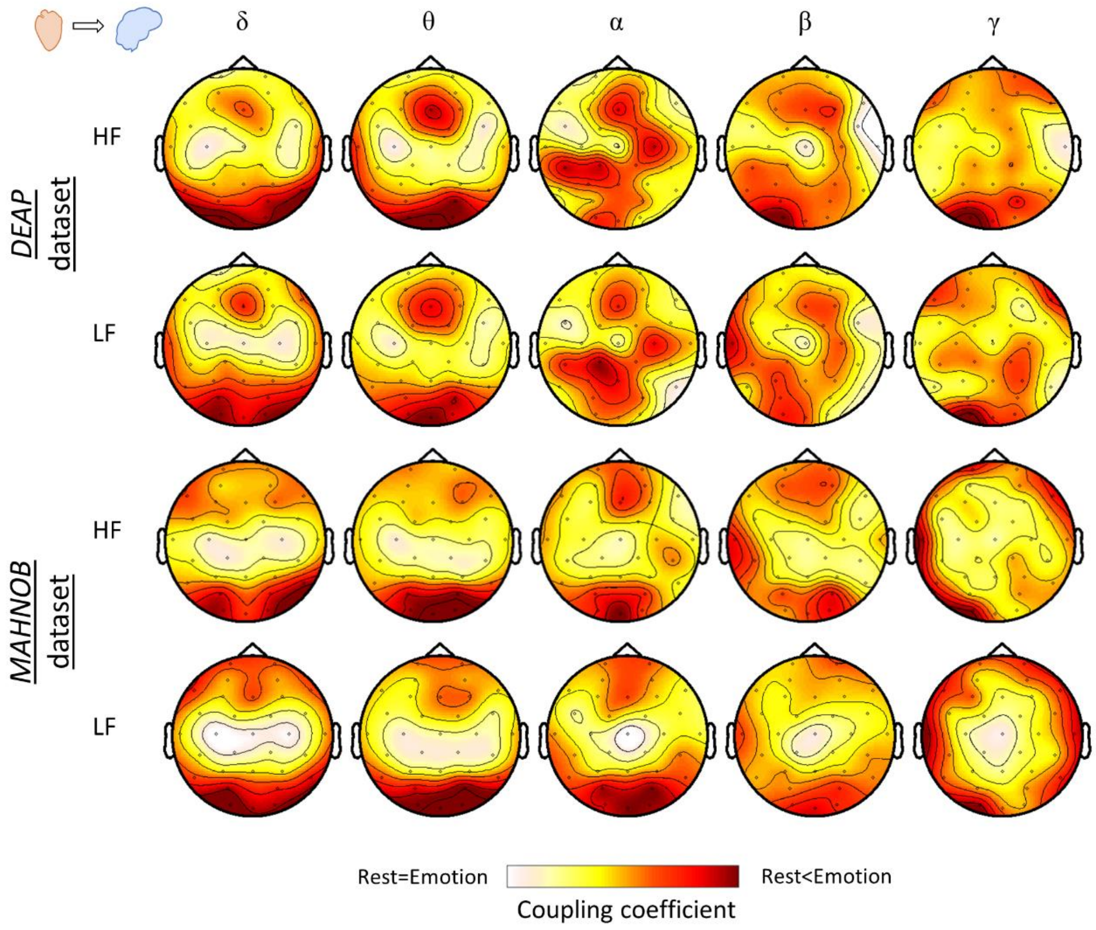


*Figure 3. Scalp regions associated with higher coupling in Heart-to-Brain Interplay. Colormaps correspond to the group-median of the time windows in which the trials present a significant change from rest, based on the cluster-based permutation analysis. Before the group-median computation, z-score was applied to the coupling coefficients among channels for individual subjects in order to highlight the regions with major activations.*

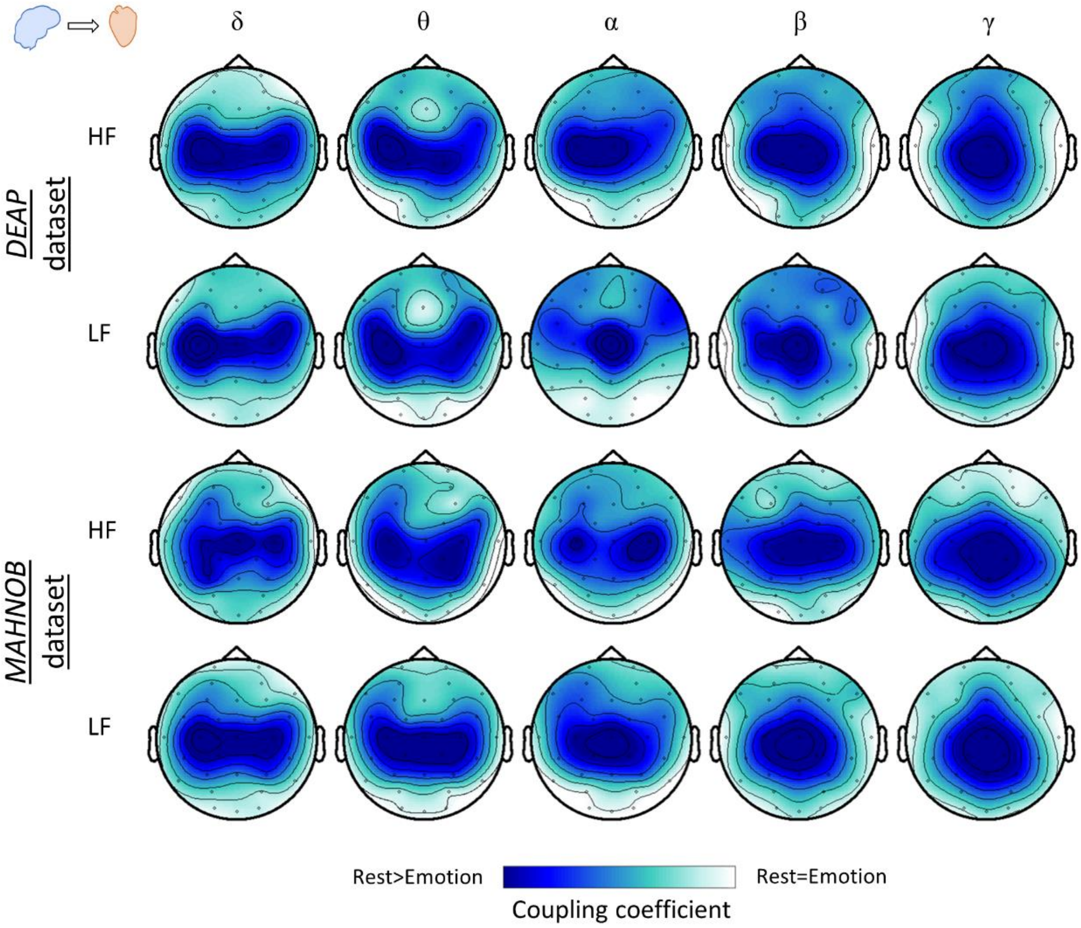


*Figure 4. Scalp regions associated with higher coupling in Brain-to-Heart Interplay. Colormaps correspond to the group-median of the time windows in which the trials present a significant change from rest, based on the cluster-based permutation analysis. Before the group-median computation, z-score was applied to the coupling coefficients among channels for individual subjects in order to highlight the regions with major activations.*

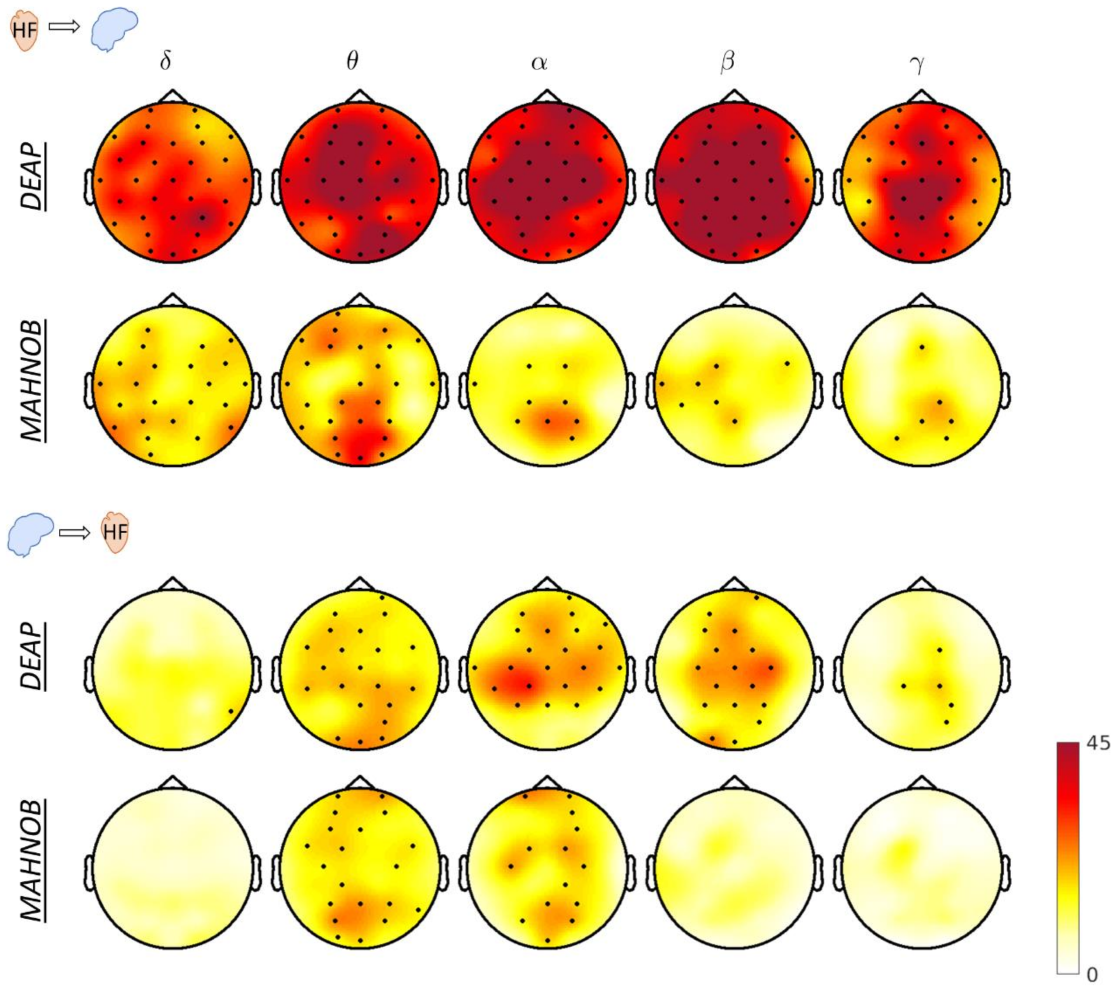


*Figure 5. Friedman test of HF-to-brain and brain-to-HF performed over four averaged time intervals: rest and first three quarters of the trial, all trials averaged separately for the two datasets. Colormaps indicate the amplitudes of the respective statistics and thick electrodes indicate significance ($p < 0.05/32$).*

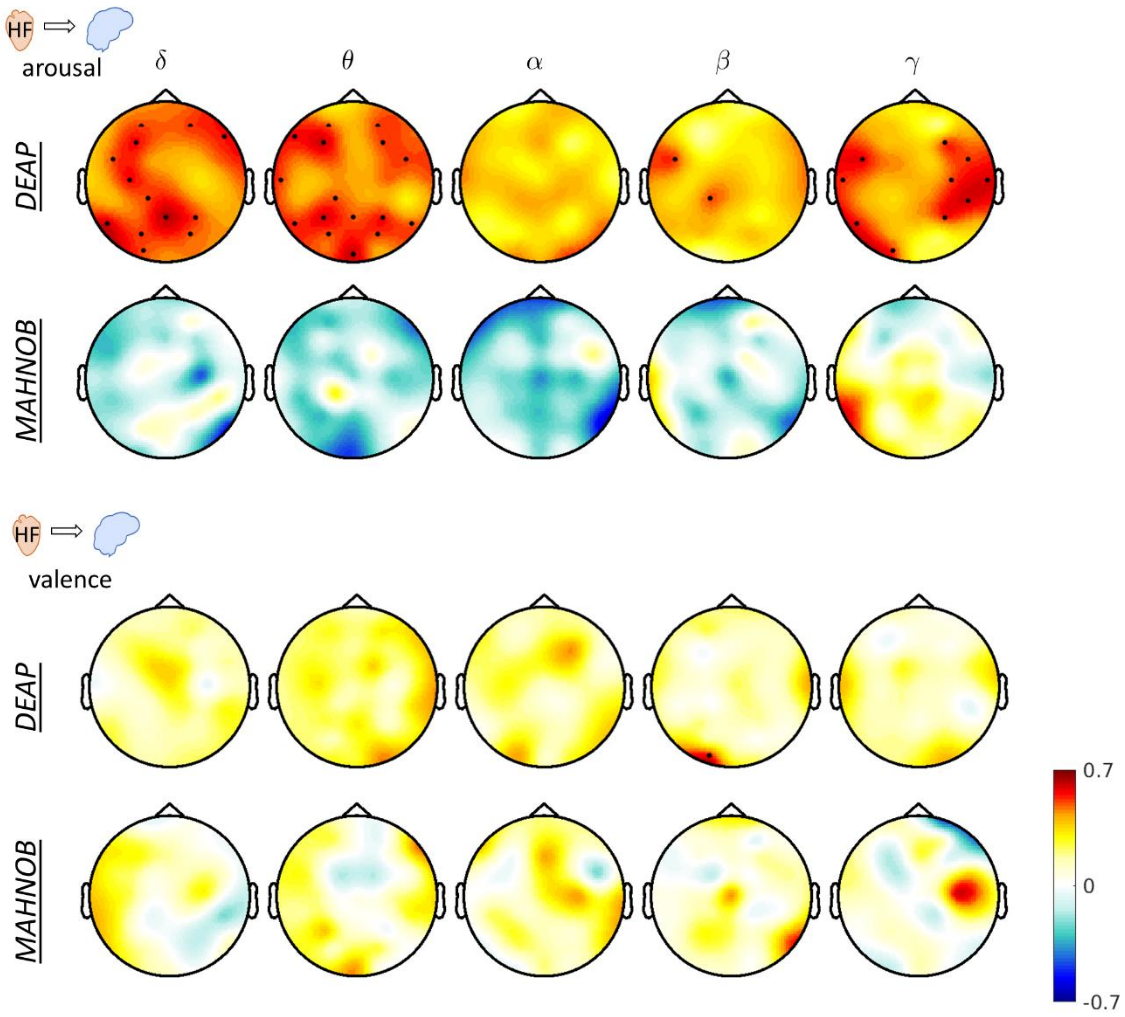


*Figure 6. Correlation analysis between ascending coupling coefficients and perceived arousal and valence. Each point evaluated in the correlation analysis corresponds to the group-median value per trial, for perceived arousal or valence, and coupling coefficients. Individual values were previously max-min normalized within the 40 trials (DEAP) or 20 trials (MAHNOB) for each subject. Colormaps indicate correlation coefficients (R), and thick electrodes indicate significant correlation corrected for multiple comparisons among channels ($p < 0.05/32$).*

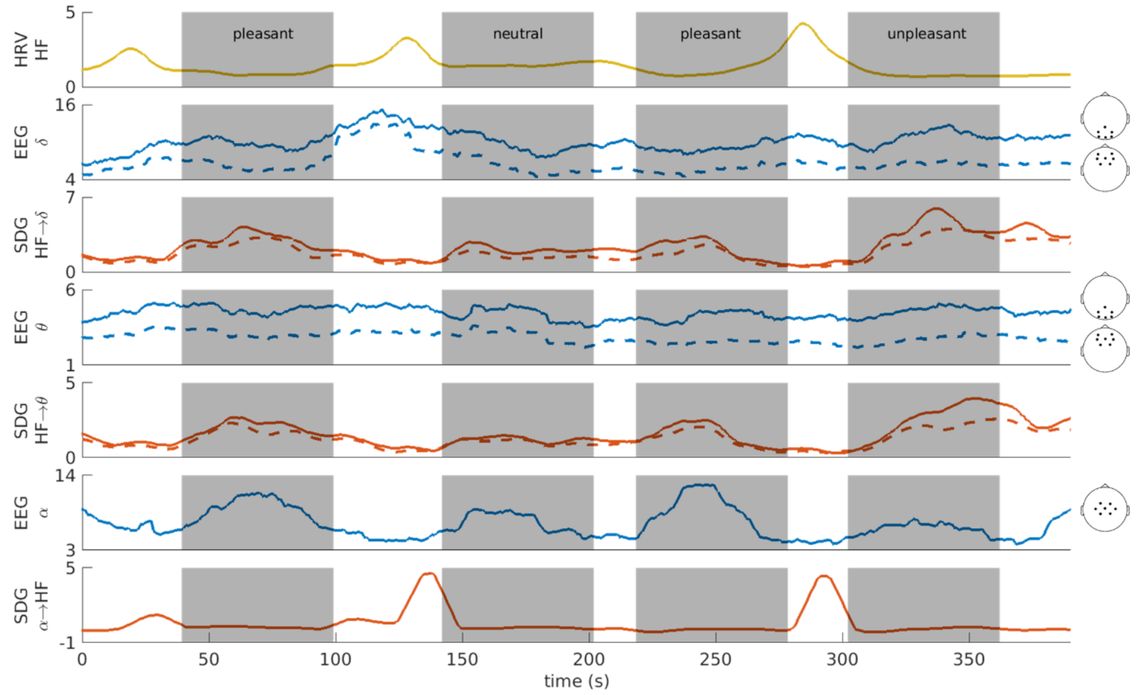


*Figure 7. Brain-Heart Interplay of an exemplary subject from the DEAP dataset. Different components of the interplay are displayed in separated panels. Shaded areas correspond to the visualization of a video trial. Components from ascending modulations were represented in two averaged groups of electrodes, centred in 'Fz' (dashed) and 'Pz' (continuous), as indicated in the figure. Components of descending modulations were represented in an averaged group of electrodes centred in 'Cz'. [2 columns]*

*Table 1. DEAP dataset trial classification. Trial numbers correspond to the original assignment in the dataset. Values reported for valence, arousal and dominance correspond to the group median.*

| *DEAP* dataset | | | | | | | | | | | | | | |
|---|---|---|---|---|---|---|---|---|---|---|---|---|---|---|
| Pleasant trials | | | | | | | | | | | | | | |
| Trial: | 1 | 2 | 3 | 4 | 5 | 6 | 7 | 8 | 9 | 11 | 14 | 18 | 19 | 20 |
| Valence: | 7 | 6.9 | 7.5 | 7 | 6.6 | 6.9 | 6.1 | 7.1 | 7.1 | 7.3 | 8 | 7.3 | 7.3 | 7 |
| Arousal: | 5.8 | 6.6 | 6.7 | 6 | 7 | 6 | 6.3 | 5.7 | 6 | 5.7 | 5 | 5 | 6.1 | 5.8 |
| Dominance: | 6.4 | 6 | 7 | 6.1 | 7 | 6.1 | 6.1 | 6.4 | 6.2 | 6.1 | 6.5 | 6 | 6.1 | 6 |
| Unpleasant or less pleasant trials | | | | | | | | | | | | | | |
| Trial: | 10 | 24 | 30 | 31 | 32 | 33 | 34 | 35 | 36 | 37 | 38 | 39 | 40 | |
| Valence: | 5 | 3.3 | 3.1 | 4 | 4 | 4.1 | 4.1 | 3.1 | 3.9 | 3 | 2.2 | 3.2 | 5 | |
| Arousal: | 6 | 5 | 6 | 6 | 7 | 5.8 | 6.4 | 6 | 6.6 | 6.1 | 6.2 | 6.2 | 5.4 | |
| Dominance: | 7.1 | 4.1 | 3.8 | 3.8 | 5.8 | 5.1 | 5.4 | 5 | 5.9 | 4 | 4.8 | 5 | 6 | |
| Neutral trials | | | | | | | | | | | | | | |
| Trial: | 12 | 13 | 15 | 16 | 17 | 21 | 22 | 23 | 25 | 26 | 27 | 28 | 29 | |
| Valence: | 6.6 | 7.2 | 6.1 | 5 | 6 | 3.6 | 4.2 | 3 | 4 | 5 | 5.3 | 4.7 | 4 | |
| Arousal: | 4 | 4.5 | 3.9 | 3.8 | 4 | 4.5 | 3.9 | 3.8 | 4 | 3.2 | 4.8 | 3.5 | 5 | |
| Dominance: | 5 | 5.1 | 5 | 4.7 | 5 | 4.8 | 4.6 | 3.3 | 4.1 | 5 | 5 | 4 | 4 | |

*Table 2. MAHNOB dataset trial classification. Trial numbers correspond to the original assignment in the dataset. Values reported for valence, arousal and dominance correspond to the group median.*

| *MAHNOB* dataset | | | | | | | | |
|---|---|---|---|---|---|---|---|---|
| Pleasant trials | | | | | | | | |
| Trial: | 3 | 5 | 7 | 9 | 10 | 15 | 18 | 19 |
| Valence: | 4 | 7 | 4 | 8 | 6 | 7 | 4 | 7 |
| Arousal: | 6 | 4 | 5 | 6 | 4 | 5 | 5.5 | 5 |
| Dominance: | 4 | 7 | 4 | 6 | 6 | 7 | 5.5 | 7 |
| Unpleasant or less pleasant trials | | | | | | | | |
| Trial: | 1 | 4 | 6 | 11 | 12 | 13 | | |
| Valence: | 2 | 1.5 | 2 | 3 | 2 | 3 | | |
| Arousal: | 6.5 | 7 | 6 | 7 | 6 | 4 | | |
| Dominance: | 3 | 2 | 4 | 4 | 3 | 4 | | |
| Neutral trials | | | | | | | | |
| Trial: | 2 | 8 | 14 | 16 | 17 | 20 | | |
| Valence: | 6 | 7 | 3 | 5 | 5 | 5 | | |
| Arousal: | 3 | 3.5 | 3 | 1.5 | 1 | 2 | | |
| Dominance: | 7 | 8 | 5 | 6.5 | 7 | 7 | | |

*Table 3. DEAP dataset video list*

| Trial | *Lastfm* tag | Artist | Title |
|---|---|---|---|
| 1 | fun | Emilíana Torrini | Jungle Drum |
| 2 | exciting | Lustra | Scotty Doesn't Know |
| 3 | joy | Jackson 5 | Blame It On The Boogie |
| 4 | | The B52'S | Love Shack |
| 5 | | Blur | Song 2 |
| 6 | | Blink 182 | First Date |
| 7 | | Benny Benassi | Satisfaction |
| 8 | | Lily Allen | F*ck You |
| 9 | | Queen | I Want To Break Free |
| 10 | | Rage Against The Machine | Bombtrack |
| 11 | happy | Michael Franti & Spearhead | Say Hey (I Love You) |
| 12 | cheerful | Grand Archives | Miniature Birds |
| 13 | love | Bright Eyes | First Day Of My Life |
| 14 | happy | Jason Mraz | I'm Yours |
| 15 | lovely | Bishop Allen | Butterfly Nets |
| 16 | sentimental | The Submarines | Darkest Things |
| 17 | | Air | Moon Safari |
| 18 | | Louis Armstrong | What A Wonderful World |
| 19 | | Manu Chao | Me Gustas Tu |
| 20 | | Taylor Swift | Love Story |
| 21 | | Diamanda Galas | Gloomy Sunday |
| 22 | sentimental | Porcupine Tree | Normal |
| 23 | melancholy | Wilco | How To Fight Loneliness |
| 24 | sad | James Blunt | Goodbye My Lover |
| 25 | depressing | A Fine Frenzy | Goodbye My Almost Lover |
| 26 | mellow | Kings Of Convenience | The Weight Of My Words |
| 27 | | Madonna | Rain |
| 28 | | Sia | Breathe Me |

| | | | |
|---|---|---|---|
| 29 | | Christina Aguilera | Hurt |
| 30 | | Enya | May It Be (Saving Private Ryan) |
| 31 | terrible | Mortemia | The One I Once Was |
| 32 | shock | Marilyn Manson | The Beautiful People |
| 33 | hate | Dead To Fall | Bastard Set Of Dreams |
| 34 | | Dj Paul Elstak | A Hardcore State Of Mind |
| 35 | | Napalm Death | Procrastination On The Empty Vessel |
| 36 | | Sepultura | Refuse Resist |
| 37 | | Cradle Of Filth | Scorched Earth Erotica |
| 38 | | Gorgoroth | Carving A Giant |
| 39 | | Dark Funeral | My Funeral |
| 40 | | Arch Enemy | My Apocalypse |

***Table 4. MAHNOB dataset video list***

| Trial | Video file name | Video description | Emotion |
|---|---|---|---|
| 1 | '30.avi' | Silent Hill | Fear |
| 2 | '52.avi' | Mr Bean's Holiday | Amusement |
| 3 | '53.avi' | Kill Bill Vol 1 | Amusement |
| 4 | '55.avi' | The Pianist | Anger/Sadness |
| 5 | '58.avi' | Mr Bean's Holiday | Amusement |
| 6 | '69.avi' | Hannibal | Disgust |
| 7 | '73.avi' | The Shining | Fear |
| 8 | '79.avi' | The Thin Red Line | Joy |
| 9 | '80.avi' | Love Actually | Joy |
| 10 | '90.avi' | Love Actually | Joy |
| 11 | '107.avi' | The Shining | Fear |
| 12 | '111.avi' | American History X | Sadness |
| 13 | '138.avi' | The Thin Red Line | Sadness |
| 14 | '146.avi' | Gangs of New York | Sadness |
| 15 | 'cats_f.avi' | | Joy |
| 16 | 'dallas_f.avi' | | Neutral |
| 17 | 'detroit_f.avi' | | Neutral |
| 18 | 'earworm_f.avi' | | Disgust |
| 19 | 'funny_f.avi' | | Joy |
| 20 | 'newyork_f.avi' | | Neutral |

***Table 5. Emotion elicitation trials from the DEAP dataset. Trials are classified as pleasant, unpleasant or neutral based on their valence and arousal self-assessment. "✓" indicates that the trial is significantly different from rest in the cluster-based permutation test (monte carlo p < 0.01).***

| *DEAP* dataset | | | | | | | | | | | | | | |
|---|---|---|---|---|---|---|---|---|---|---|---|---|---|---|
| Pleasant trials | | | | | | | | | | | | | | |
| Trial: | 1 | 2 | 3 | 4 | 5 | 6 | 7 | 8 | 9 | 11 | 14 | 18 | 19 | 20 | Total |
| LF → Brain | ✓ | | ✓ | ✓ | ✓ | ✓ | ✓ | | ✓ | ✓ | ✓ | | | ✓ | 10/14 |
| HF → Brain | ✓ | ✓ | ✓ | ✓ | ✓ | ✓ | ✓ | | ✓ | ✓ | ✓ | ✓ | ✓ | ✓ | 13/14 |

| | | | | | | | | | | | | | | | |
|---|---|---|---|---|---|---|---|---|---|---|---|---|---|---|---|
| Brain → LF | | ✓ | | | ✓ | ✓ | ✓ | | ✓ | | | | | | 5/14 |
| Brain → HF | ✓ | ✓ | ✓ | ✓ | ✓ | ✓ | ✓ | ✓ | ✓ | ✓ | ✓ | ✓ | ✓ | ✓ | 14/14 |
| Unpleasant or less pleasant trials | | | | | | | | | | | | | | | |
| Trial: | 10 | 24 | 30 | 31 | 32 | 33 | 34 | 35 | 36 | 37 | 38 | 39 | 40 | Total | |
| LF → Brain | | ✓ | ✓ | | | ✓ | | ✓ | ✓ | ✓ | | ✓ | ✓ | 8/13 | |
| HF → Brain | ✓ | ✓ | ✓ | | ✓ | ✓ | | ✓ | ✓ | ✓ | ✓ | ✓ | ✓ | 11/13 | |
| Brain → LF | | | | | ✓ | | | ✓ | | | | ✓ | | 3/13 | |
| Brain → HF | ✓ | ✓ | ✓ | | ✓ | ✓ | ✓ | ✓ | ✓ | ✓ | ✓ | ✓ | ✓ | 12/13 | |
| Neutral trials | | | | | | | | | | | | | | | |
| Trial: | 12 | 13 | 15 | 16 | 17 | 21 | 22 | 23 | 25 | 26 | 27 | 28 | 29 | Total | |
| LF → Brain | ✓ | | ✓ | ✓ | ✓ | ✓ | ✓ | | ✓ | | | | ✓ | 7/13 | |
| HF → Brain | ✓ | ✓ | ✓ | ✓ | ✓ | ✓ | ✓ | | ✓ | ✓ | ✓ | | ✓ | 11/13 | |
| Brain → LF | | | | | | | | | | | | | | 0/13 | |
| Brain → HF | ✓ | ✓ | ✓ | ✓ | ✓ | ✓ | ✓ | ✓ | ✓ | ✓ | | ✓ | ✓ | 12/13 | |

***Table 6. Emotion elicitation trials from the MAHNOB dataset. Trials are classified as pleasant, unpleasant or neutral based on their valence and arousal self-assessment. "✓" indicates that the trial is significantly different from rest in the cluster-based permutation test (monte carlo p < 0.01).***

| *MAHNOB* dataset | | | | | | | | | |
|---|---|---|---|---|---|---|---|---|---|
| Pleasant trials | | | | | | | | | |
| Trial: | 3 | 5 | 7 | 9 | 10 | 15 | 18 | 19 | Total |
| LF → Brain | ✓ | ✓ | | ✓ | | | | ✓ | 4/8 |
| HF → Brain | ✓ | ✓ | | ✓ | | | | | 3/8 |
| Brain → LF | | ✓ | | | | | | | 1/8 |
| Brain → HF | | | | ✓ | | ✓ | ✓ | ✓ | 4/8 |
| Unpleasant or less pleasant trials | | | | | | | | | |
| Trial: | 1 | 4 | 6 | 11 | 12 | 13 | Total | | |
| LF → Brain | | ✓ | ✓ | | ✓ | ✓ | 4/6 | | |
| HF → Brain | | ✓ | ✓ | | ✓ | ✓ | 4/6 | | |
| Brain → LF | ✓ | ✓ | | | ✓ | | 3/6 | | |
| Brain → HF | | | | | ✓ | | 1/6 | | |
| Neutral trials | | | | | | | | | |
| Trial: | 2 | 8 | 14 | 16 | 17 | 20 | Total | | |
| LF → Brain | ✓ | | ✓ | ✓ | ✓ | ✓ | 5/6 | | |
| HF → Brain | ✓ | ✓ | | ✓ | ✓ | ✓ | 5/6 | | |
| Brain → LF | | | | | | | 0/6 | | |
| Brain → HF | ✓ | | ✓ | ✓ | | | 3/6 | | |

***Table 7. Cluster-based permutation statistics in the DEAP dataset HF-Brain interplay. The latency indicates the time window in which the cluster analysis found significant changes. Z-value is the statistic from the paired Wilcoxon test performed between emotion elicitation and resting state, averaging all data points defined in the cluster analysis. P-value is the p associated to the Z-value and Montecarlo P-value is computed out of 10000 random partitions of the data.***

| | | HF → Brain | | | | Brain → HF | | | |
|---|---|---|---|---|---|---|---|---|---|
| | trial | latency (s) | z | p | monte carlo p | latency (s) | z | p | monte carlo p |
| [illegible] | 1 | 5-52 | 3.63 | 0.0003 | <0.0001 | 9-58 | -3.40 | 0.0007 | 0.0002 |

| | | | | | | | | | |
|---|---|---|---|---|---|---|---|---|---|
| | 2 | 0-49 | 3.61 | 0.0003 | 0.0002 | 1-59 | -3.35 | 0.0008 | 0.0005 |
| | 3 | 2-57 | 3.68 | 0.0002 | 0.0003 | 0-59 | -3.05 | 0.0023 | 0.0012 |
| | 4 | 2-59 | 4.02 | <0.0001 | 0.0001 | 14-59 | -3.23 | 0.0012 | 0.0008 |
| | 5 | 8-55 | 4.13 | <0.0001 | <0.0001 | 16-44 | -3.38 | 0.0007 | 0.0003 |
| | 6 | 0-55 | 3.68 | 0.0002 | 0.0001 | 11-59 | -3.55 | 0.0004 | 0.0002 |
| | 7 | 12-57 | 3.48 | 0.0005 | 0.0002 | 10-59 | -3.81 | 0.0001 | <0.0001 |
| | 8 | - | - | - | - | 16-50 | -2.92 | 0.0035 | 0.0021 |
| | 9 | 19-59 | 3.48 | 0.0005 | 0.0003 | 13-59 | -4.24 | <0.0001 | <0.0001 |
| | 11 | 6-50 | 3.93 | <0.0001 | <0.0001 | 10-59 | -3.44 | 0.0006 | 0.0001 |
| | 14 | 13-54 | 4.41 | <0.0001 | <0.0001 | 0-59 | -3.43 | 0.0006 | 0.0004 |
| | 18 | 0-50 | 4.28 | <0.0001 | <0.0001 | 0-59 | -4.23 | <0.0001 | <0.0001 |
| | 19 | 7-54 | 3.98 | <0.0001 | <0.0001 | 14-59 | -3.35 | 0.0008 | 0.0003 |
| | 20 | 0-56 | 4.36 | <0.0001 | <0.0001 | 6-59 | -4.56 | <0.0001 | <0.0001 |
| unpleasant | 10 | 14-59 | 3.42 | 0.0006 | 0.0003 | 12-59 | -2.80 | 0.0050 | 0.0040 |
| | 24 | 14-53 | 3.55 | 0.0004 | 0.0001 | 3-56 | -3.46 | 0.0005 | 0.0001 |
| | 30 | 12-47 | 3.68 | 0.0002 | <0.0001 | 12-59 | -3.52 | 0.0004 | 0.0002 |
| | 31 | 2-59 | 3.96 | <0.0001 | <0.0001 | 22-59 | -3.17 | 0.0015 | 0.0005 |
| | 32 | 12-59 | 4.00 | <0.0001 | 0.0001 | 40-59 | -3.35 | 0.0008 | 0.0006 |
| | 33 | 22-45 | 3.46 | 0.0005 | 0.0002 | 0-42 | -3.30 | 0.0010 | 0.0003 |
| | 34 | 25-39 | 3.31 | 0.0009 | 0.0008 | 0-59 | -3.21 | 0.0013 | 0.0004 |
| | 35 | 10-56 | 4.45 | <0.0001 | <0.0001 | 12-46 | -3.57 | 0.0004 | 0.0001 |
| | 36 | 15-59 | 4.28 | <0.0001 | <0.0001 | 15-59 | -3.25 | 0.0011 | 0.0013 |
| | 37 | 16-53 | 3.93 | <0.0001 | <0.0001 | 13-53 | -3.40 | 0.0007 | 0.0001 |
| | 38 | 18-59 | 3.53 | 0.0004 | 0.0001 | 13-59 | -4.10 | <0.0001 | <0.0001 |
| | 39 | 0-54 | 4.15 | <0.0001 | <0.0001 | 0-41 | -3.95 | <0.0001 | <0.0001 |
| | 40 | 10-59 | 3.76 | 0.0002 | 0.0001 | 12-59 | -3.42 | 0.0006 | 0.0006 |
| neutral | 12 | 6-57 | 3.55 | 0.0004 | 0.0001 | 11-59 | -3.89 | 0.0001 | <0.0001 |
| | 13 | 37-52 | 3.29 | 0.0010 | 0.0009 | 13-59 | -3.91 | <0.0001 | <0.0001 |
| | 15 | 12-44 | 3.52 | 0.0004 | 0.0003 | 15-57 | -2.92 | 0.0035 | 0.0028 |
| | 16 | 19-59 | 3.61 | 0.0003 | <0.0001 | 13-59 | -3.55 | 0.0004 | 0.0002 |
| | 17 | 36-53 | 3.38 | 0.0007 | 0.0003 | 11-59 | -2.94 | 0.0033 | 0.0017 |
| | 21 | 20-58 | 3.74 | 0.0002 | <0.0001 | 23-59 | -3.35 | 0.0008 | 0.0006 |
| | 22 | 0-53 | 4.04 | <0.0001 | <0.0001 | 14-59 | -3.40 | 0.0007 | 0.0004 |
| | 23 | - | - | - | - | 9-58 | -2.84 | 0.0045 | 0.0027 |
| | 25 | 16-18 | 3.07 | 0.0022 | 0.0013 | 5-59 | -2.84 | 0.0045 | 0.0032 |
| | 26 | 14-56 | 3.63 | 0.0003 | 0.0002 | 13-58 | -3.57 | 0.0004 | 0.0001 |
| | 27 | 12-38 | 4.24 | <0.0001 | <0.0001 | 13-42 | -3.25 | 0.0011 | 0.0006 |
| | 28 | - | - | - | - | 16-59 | -3.57 | 0.0004 | 0.0001 |
| | 29 | 0-55 | 4.17 | <0.0001 | <0.0001 | 0-59 | -3.44 | 0.0006 | 0.0003 |

*Table 8. Cluster-based permutation statistics in the DEAP dataset LF-Brain interplay. The latency indicates the time window in which the cluster analysis found significant changes. Z-value is the statistic from the paired Wilcoxon test performed between emotion elicitation and resting state, averaging all data points defined in the cluster*

***analysis. P-value is the p associated to the Z-value and Montecarlo P-value is computed out of 10000 random partitions of the data.***

| | | LF → Brain | | | | Brain → LF | | | |
|---|---|---|---|---|---|---|---|---|---|
| | trial | latency (s) | z | p | monte carlo p | latency (s) | z | p | monte carlo p |
| pleasant | 1 | 26-45 | 3.48 | 0.0005 | 0.0003 | - | - | - | - |
| | 2 | - | - | - | - | - | - | - | - |
| | 3 | 26-38 | 3.08 | 0.0021 | 0.0013 | - | - | - | - |
| | 4 | 21-50 | 3.27 | 0.0011 | 0.0005 | - | - | - | - |
| | 5 | 27-45 | 3.31 | 0.0009 | 0.0003 | 37-42 | -2.86 | 0.0042 | 0.0032 |
| | 6 | 12-51 | 3.48 | 0.0005 | 0.0001 | 22-59 | -3.37 | 0.0008 | 0.0003 |
| | 7 | 25-47 | 3.05 | 0.0023 | 0.0022 | 47-59 | -3.18 | 0.0015 | 0.0009 |
| | 8 | - | - | - | - | - | - | - | - |
| | 9 | 25-41 | 3.22 | 0.0013 | 0.0009 | 34-41 | -2.82 | 0.0047 | 0.0030 |
| | 11 | 27-38 | 3.10 | 0.0019 | 0.0015 | - | - | - | - |
| | 14 | 23-50 | 3.59 | 0.0003 | <0.0001 | - | - | - | - |
| | 18 | - | - | - | - | - | - | - | - |
| | 19 | - | - | - | - | - | - | - | - |
| | 20 | 22-34 | 3.20 | 0.0014 | 0.0014 | - | - | - | - |
| unpleasant | 10 | - | - | - | - | - | - | - | - |
| | 24 | 17-39 | 3.22 | 0.0013 | 0.0003 | - | - | - | - |
| | 30 | 17-42 | 3.44 | 0.0006 | 0.0003 | - | - | - | - |
| | 31 | 36-49 | 3.33 | 0.0009 | 0.0008 | - | - | - | - |
| | 32 | 33-36 | 3.22 | 0.0013 | 0.0006 | 25-36 | -3.03 | 0.0025 | 0.0017 |
| | 33 | 31-35 | 2.97 | 0.0029 | 0.0020 | - | - | - | - |
| | 34 | - | - | - | - | - | - | - | - |
| | 35 | 20-53 | 3.95 | <0.0001 | <0.0001 | 49-54 | -2.71 | 0.0067 | 0.0063 |
| | 36 | 28-40 | 3.29 | 0.0010 | 0.0004 | - | - | - | - |
| | 37 | 25-47 | 3.33 | 0.0009 | 0.0006 | - | - | - | - |
| | 38 | - | - | - | - | - | - | - | - |
| | 39 | 23-52 | 3.65 | 0.0003 | 0.0002 | - | - | - | - |
| | 40 | 36-49 | 3.25 | 0.0011 | 0.0007 | - | - | - | - |
| neutral | 12 | 19-43 | 3.46 | 0.0005 | 0.0003 | - | - | - | - |
| | 13 | - | - | - | - | - | - | - | - |
| | 15 | 27-31 | 2.79 | 0.0053 | 0.0039 | - | - | - | - |
| | 16 | 25-26 | -0.00 | 1.0000 | 1.0000 | - | - | - | - |
| | 17 | 36-50 | 3.18 | 0.0015 | 0.0008 | - | - | - | - |
| | 21 | 34-44 | 3.07 | 0.0022 | 0.0013 | - | - | - | - |
| | 22 | 14-18 | 3.07 | 0.0022 | 0.0010 | - | - | - | - |
| | 23 | - | - | - | - | - | - | - | - |
| | 25 | 36-41 | 3.38 | 0.0007 | 0.0003 | - | - | - | - |
| | 26 | - | - | - | - | - | - | - | - |
| | 27 | - | - | - | - | - | - | - | - |
| | 28 | - | - | - | - | - | - | - | - |
| | 29 | 17-48 | 3.55 | 0.0004 | 0.0003 | - | - | - | - |

*Table 9. Cluster-based permutation statistics in the MAHNOB dataset HF-Brain interplay. The latency indicates the time window in which the cluster analysis found significant changes. Z-value is the statistic from the paired Wilcoxon test performed between emotion elicitation and resting state, averaging all data points defined in the cluster analysis. P-value is the p associated to the Z-value and Montecarlo P-value is computed out of 10000 random partitions of the data.*

| | | HF ⟶ Brain | | | | Brain ⟶ HF | | | |
|---|---|---|---|---|---|---|---|---|---|
| | trial | latency (s) | z | p | monte carlo p | latency (s) | z | p | monte carlo p |
| pleasant | 3 | 16-69 | 3.35 | 0.0008 | 0.0003 | - | - | - | - |
| | 5 | 9-41 | 3.95 | <0.0001 | <0.0001 | - | - | - | - |
| | 7 | - | - | - | - | - | - | - | - |
| | 9 | 10-94 | 3.22 | 0.0013 | 0.0006 | 81-94 | -3.27 | 0.0011 | 0.0006 |
| | 10 | - | - | - | - | - | - | - | - |
| | 15 | - | - | - | - | 62-92 | -3.27 | 0.0011 | 0.0003 |
| | 18 | - | - | - | - | 27-53 | -3.19 | 0.0014 | 0.0006 |
| | 19 | - | - | - | - | 53-71 | -3.15 | 0.0016 | 0.0011 |
| unpleasant | 1 | - | - | - | - | - | - | - | - |
| | 4 | 17-44 | 3.54 | 0.0004 | 0.0003 | - | - | - | - |
| | 6 | 10-38 | 3.24 | 0.0012 | 0.0003 | - | - | - | - |
| | 11 | - | - | - | - | - | - | - | - |
| | 12 | 39-100 | 3.32 | 0.0009 | 0.0005 | 24-80 | -2.97 | 0.0029 | 0.0021 |
| | 13 | 4-101 | 3.70 | 0.0002 | 0.0001 | - | - | - | - |
| neutral | 2 | 4-71 | 4.35 | <0.0001 | <0.0001 | 26-45 | -3.15 | 0.0016 | 0.0007 |
| | 8 | 11-21 | 3.57 | 0.0004 | 0.0001 | - | - | - | - |
| | 14 | - | - | - | - | 32-87 | -3.10 | 0.0019 | 0.0011 |
| | 16 | 35-79 | 3.47 | 0.0005 | 0.0002 | 34-66 | -3.47 | 0.0005 | 0.0005 |
| | 17 | 46-72 | 3.22 | 0.0013 | 0.0006 | 85-88 | -0.00 | 1.0000 | 1.0000 |
| | 20 | 41-62 | 3.30 | 0.0010 | 0.0003 | - | - | - | - |

*Table 10. Cluster-based permutation statistics in the MAHNOB dataset LF-Brain interplay. The latency indicates the time window in which the cluster analysis found significant changes. Z-value is the statistic from the paired Wilcoxon test performed between emotion elicitation and resting state, averaging all data points defined in the cluster analysis. P-value is the p associated to the Z-value and Montecarlo P-value is computed out of 10000 random partitions of the data.*

| | | LF ⟶ Brain | | | | Brain ⟶ LF | | | |
|---|---|---|---|---|---|---|---|---|---|
| | trial | latency (s) | z | p | monte carlo p | latency (s) | z | p | monte carlo p |
| pleasant | 3 | 41-98 | 3.70 | 0.0002 | 0.0001 | - | - | - | - |
| | 5 | 22-42 | 3.42 | 0.0006 | 0.0001 | 28-58 | -2.96 | 0.0031 | 0.0019 |
| | 7 | - | - | - | - | - | - | - | - |
| | 9 | 23-88 | 3.22 | 0.0013 | 0.0003 | - | - | - | - |
| | 10 | - | - | - | - | - | - | - | - |
| | 15 | - | - | - | - | - | - | - | - |

| | | | | | | | | | |
|---|---|---|---|---|---|---|---|---|---|
| | 18 | - | - | - | - | - | - | - | - |
| | 19 | - | - | - | - | - | - | - | - |
| unpleasant | 1 | - | - | - | - | 62-70 | -2.70 | 0.0068 | 0.0061 |
| | 4 | 12-55 | 3.42 | 0.0006 | 0.0002 | 13-76 | -3.52 | 0.0004 | 0.0004 |
| | 6 | 11-58 | 3.60 | 0.0003 | 0.0001 | - | - | - | - |
| | 11 | - | - | - | - | - | - | - | - |
| | 12 | 25-101 | 3.57 | 0.0004 | <0.0001 | 13-113 | -3.54 | 0.0004 | 0.0001 |
| | 13 | 12-102 | 3.80 | 0.0001 | <0.0001 | - | - | - | - |
| neutral | 2 | 10-26 | 2.98 | 0.0029 | 0.0024 | - | - | - | - |
| | 8 | - | - | - | - | - | - | - | - |
| | 14 | 67-74 | 3.00 | 0.0027 | 0.0018 | - | - | - | - |
| | 16 | 38-88 | 3.47 | 0.0005 | 0.0002 | - | - | - | - |
| | 17 | 52-64 | 3.10 | 0.0019 | 0.0010 | - | - | - | - |
| | 20 | 18-88 | 3.51 | 0.0004 | 0.0002 | - | - | - | - |
